# SmERF6 promotes the expression of terpenoid pathway in Salvia officinalis and improves the production of high value abietane diterpenes, carnosol and carnosic acid

**DOI:** 10.1101/2023.11.23.568411

**Authors:** Revuru Bharadwaj, Gayathri Thashanamoorthi, Pratibha Demiwal, Debabrata Sircar, Sathishkumar Ramalingam

**Affiliations:** Plant Genetic Engineering Laboratory, Department of Biotechnology, Bharathiar University, Coimbatore, India; Plant Molecular Biology Laboratory, Department of Biosciences and Bioengineering, Indian Institute of Technology, Roorkee, India

**Keywords:** *Salvia officinalis*, Carnosol, Carnosic acid, Metabolic engineering, Transcription factors, Pharmaceutically important diterpenes

## Abstract

Carnosol (CO) and carnosic acid (CA) are pharmaceutically important diterpenes predominantly produced in members of Lamiaceae, *Salvia officinalis*, *Salvia fruticosa* and *Rosmarinus officinalis*. Nevertheless, availability of these compounds in plant system is very low.

With an effort to improve the *in planta* content of these diterpenes, *SmERF6* (*Salvia miltiorrhiza Ethylene Responsive Factor 6*) transcription factor was expressed in *S. officinalis* heterologously. SmERF6 is known to bind at the promoter regions of *Copalyl diphosphate synthase* (*CPS*) and *Kaurene synthase like* (*KSL*) genes and improve ferruginol content, a common precursor for abietane diterpenes in *Salvia* genus.

Transient expression of *SmERF6* exhibited the inter-specific activity in promoting differential accumulation of diterpenes in *S. officinalis*. Overexpression studies showed elevation in the levels of CO (10-folds) and CA (8-folds). Further, in infiltrated leaves levels of ferruginol (50%) and CA derivatives (rosmanol, epirosmanal, methyl carnosic acid) were significantly upregulated along with the other signature terpenes. While, knockdown of homologous *ERF6* resulted in drastic reduction of the metabolite content.

Finally, stable transgenic lines of *S. officinalis* developed through *in planta Agrobacterium* mediated genetic transformation method accumulated higher levels of CO (4-folds) and CA (3-folds) as compared to wild plants.

Overall, the present study is the first report on improving the content of pharmaceutically important diterpenes in *S. officinalis* by overexpressing pathway specific transcription factor. The current results showed convincing evidence for the concept of improving the content of specialized metabolite(s) in medicinal plants by manipulating the expression of key transcription factors.

## Introduction

Terpenes constitute the largest class of plant metabolites synthesized from 5-carbon (C_5_) building blocks, isopentenyl pyrophosphate (IPP) and dimethylallyl pyrophosphate (DMAPP). Terpenes actively participate in primary metabolism by contributing towards the production of phytohormones, accessory pigments and components of electron transfer system. In addition, few terpenes are known to act as volatiles to attract pollinators and defence compounds to deter pathogens (Pichersky and Raguso, 2018). IPP and DMAPP are the final products of mevalonic acid pathway (MVA pathway) and 2-*C*-Methyl-D-erythritol 4-phosphate pathway (MEP pathway; Nagegowda and Gupta, 2020). Head to tail condensation of IPP and DMAPP by *trans*-prenyl transferases (TPTs) leads to the formation of geranyl pyrophosphate (GPP; C_10_), farnesyl pyrophosphate (FPP; C_15_) and geranylgeranyl pyrophosphate (GGPP; C_20_), which are the central intermediaries for terpene diversity (Zhou and Pichersky, 2020a). These linear prenoid moieties are cyclized by the catalytic activity of terpene synthases (TPS) resulting in the biosynthesis of mono-, di-, sester-and tri-terpene skeletons (Chen *et al*., 2011; Zhou and Pichersky, 2020b). Subsequently, these skeletons were decorated with different functional groups by enzymes such as cytochrome P450s (CYP450s), dehydrogenases, methyl and acyl transferases resulting in plant terpene diversity (Zhou and Pichersky, 2020 a, b).

Diterpenes (C_20_) are known to be the most diversified subclass of plant terpenes amounting to >18,000 compounds. They are involved in the regulating plant growth, participating in defence also possess economic importance in pharmaceutical and agriculture sectors (Hu *et al*., 2021). The head to tail condensation of four isoprene units by GGPP synthase (GGPPS) results in the formation of GGPP. Class II diterpene synthases (diTPS eg: *ent*-copalyl diphosphate synthase) accept GGPP as substrate and perform protonation-initiated cyclization to form intermediates like copalyl diphosphate. Further, these intermediates undergo ionization of the phosphate group and subsequent cyclization by class I diTPS (eg: Kaurene synthase; Bathe and Tissier, 2019). Predominantly, diterpenes are biosynthesized through subsequent action of class II and class I diTPS. Gibberellin biosynthesis is a typical example of the process which is found to be conserved among the vascular plants (Su *et al*., 2016). The above mentioned bi-sequential reaction leads to the formation of a structurally similar group of bicyclic metabolites. Hence, the derived diterpenes are named as labdane-related diterpenes (Peters, 2010). However, few diterpenes violate the above rule and are produced through catalysis of class I diTPS. They have a macrocyclic ring and hence termed as non labdane-related diterpenes, eg: taxadiene, casbene, neocembrene (Hu *et al*., 2021). Diterpenes, such as, gibberellins and phytol are known to be constitutively produced in plant system whereas a few specialized diterpenes are produced during specific developmental stages. Interestingly, diterpenes such as, taxadiene, andrographolides, steviol glycosides, forskolin, tanshinones, CO, CA, salviol, sugiol, sclareol, casbene, neocembrene, ginkgolides and phytoalexins are the typical examples having significant commercial value in food, agriculture, cosmetic and pharmaceutical sectors (Hu *et al*., 2021; Chacón-Morales, 2022).

CO and CA are phenolic, abietane type tricyclic diterpenoid compounds, predominantly produced by members of the Lamiaceae (mint family) that include *S. officinalis* (garden sage), *S. fruticosa* and *R. officinalis* (rosemary) (Božić *et al*., 2015). CO and CA are recommended as alternative drugs to treat neurological disorders by international agencies and possess high market value (Andrade *et al*., 2018). Linde discovered CA in *S. officinalis* for the first time and Wenkert discovered the same in *R. officinalis* (Bahri *et al*., 2016). CO biosynthesis occurs in glandular trichomes of younger leaves and gets initiated with GGPP cyclization to copalyl diphosphate (CP) by Copalyl diphosphate synthase (CPS; class II diTPS) and CP is converted to miltiradiene by kaurene synthase like (KSL), a class I diTPS (Brückner *et al*., 2014; Božić *et al*., 2015). Isoforms of *CPS* and *KSL* genes were characterized in *S. fruticosa* (*SfCPS*, *SfKSL*), *R. officinalis* (*RoCPS*, *RoKSL1/RoKSL2*) and *S. officinalis* (*SoCPS1/CPS2*, *SoKSL1/KSL2*) (Brückner *et al*., 2014; Božić *et al*., 2015; Li *et al*., 2022; **Figure 1**). Subsequently, CYP450s perform scaffold rearrangement of miltiradiene to form CA. Towards this end, Scheler *et al*. (2016) characterized three bi-functional CYP450s (CYP76AH22, CYP76AH23, CYP76AH24) from *S. fruticosa* and *R. officinalis*, which convert miltiradiene to ferruginol and consequent conversion of ferruginol to 11-hydroxy ferruginol by hydroxylation at the 11^th^ position (Božić *et al*., 2015; Scheler *et al*., 2016; **Figure 1**). Further, three CYP450s namely, CYP76AK6, CYP76AK7 and CYP76AK8, were functionally characterized in *S. fruticosa* and *R. officinalis* respectively. These three CYP450s were known to play significant role in the conversion of 11-hydroxy ferruginol to CA. It is also known that the above mentioned conversion involves a multi-step process with the formation of metabolic intermediates, such as, 11,20-dihydroxyferruginol and carnosaldehyde (Scheler *et al*., 2016). Finally, CA is converted spontaneously to CO by oxidation (Scheler *et al*., 2016). In the recent times, global demand for CO and CA is escalating exponentially owing to their pharmacological significance. However, *in planta* low level accumulation of the desirable compounds is found to be a bottleneck for commercial exploitation. In addition, relatively poor multi-omics knowledge about CO accumulator species poses limitations in the in-depth understanding of pathway dynamics at chromatin level and respective regulatory mechanisms in order to produce the cherishable compounds at larger scale.

**Figure 1:**
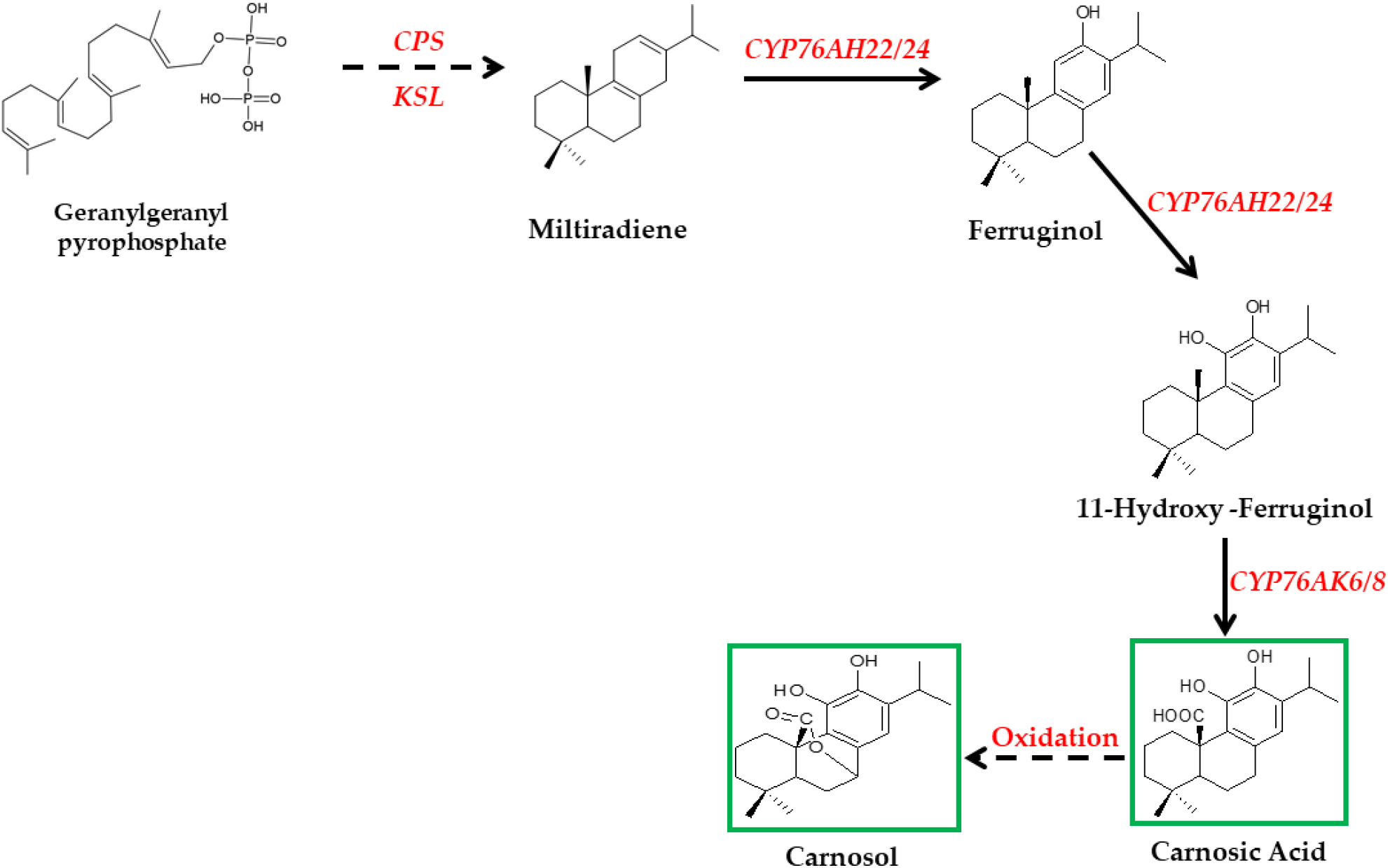
Representative biosynthetic pathway of CO in *S. officinalis*. Initially, GGPP is converted to militradiene through enzymatic steps mediated by Copalyl diphosphate synthase (CPS) and Kaurene synthase like (KSL). Later, CYP76AH22/24 converts miltiradiene to ferruginol followed by further conversion to 11-hydroxy-ferruginol. Finally, CYP76AK6/8 mediated catalysis converts 11-hydroxy-ferruginol to CA and CA is immediately converted to CO.

*Salvia miltiorrhiza* (Danshen red plant; native to China) naturally accumulates tanshinones, a group of diterpenes in the roots which are known to have pharmaceutical properties in providing protection against cardio-vascular diseases. Interestingly, tanshinones and CO are derivatives of ferruginol and share common biosynthetic pathway till the formation of ferruginol (Hu *et al*., 2021; **Figure 2**). Several reports indicate the achievement of tanshinones at significant titre in transgenic hairy roots and cell suspension systems by overexpressing pathway specific transcription factors. Towards this end, various families of transcription factors (TFs) were functionally characterized which upregulate transcription of the pathway specific genes. For example, two WRKY class TFs (*Sm*WRKY2, *Sm*WRKY34) are reported to enhance the transcription of *SmGGPPS* and *SmCPS* genes (Deng *et al*., 2019; Shi *et al*., 2022). Similarly, bHLH (basic loop helix) TFs (*Sm*bHLH10, *Sm*bHLH148) target the *DXS* and *CPS* genes in tanshinone pathway and upregulate their expression (Xing *et al*., 2018 a, b). On the other hand, GRAS and MYB (myeloblastosis) class TFs were found to have a possible role in escalating tanshinone production (Wu *et al*., 2021). Finally, three ERF (Ethylene Responsive Factor) TFs (*Sm*ERF6, *Sm*ERF8, *Sm*ERF128) were shown to bind at upstream regions of *SmCPS*, *SmKSL* and *CYP76AH1* genes in a coordinated fashion resulting in the improved bioproduction and the subsequent accumulation of the desired metabolite(s) (Bai *et al*., 2018, 2020; Zhang *et al*., 2019).

**Figure 2:**
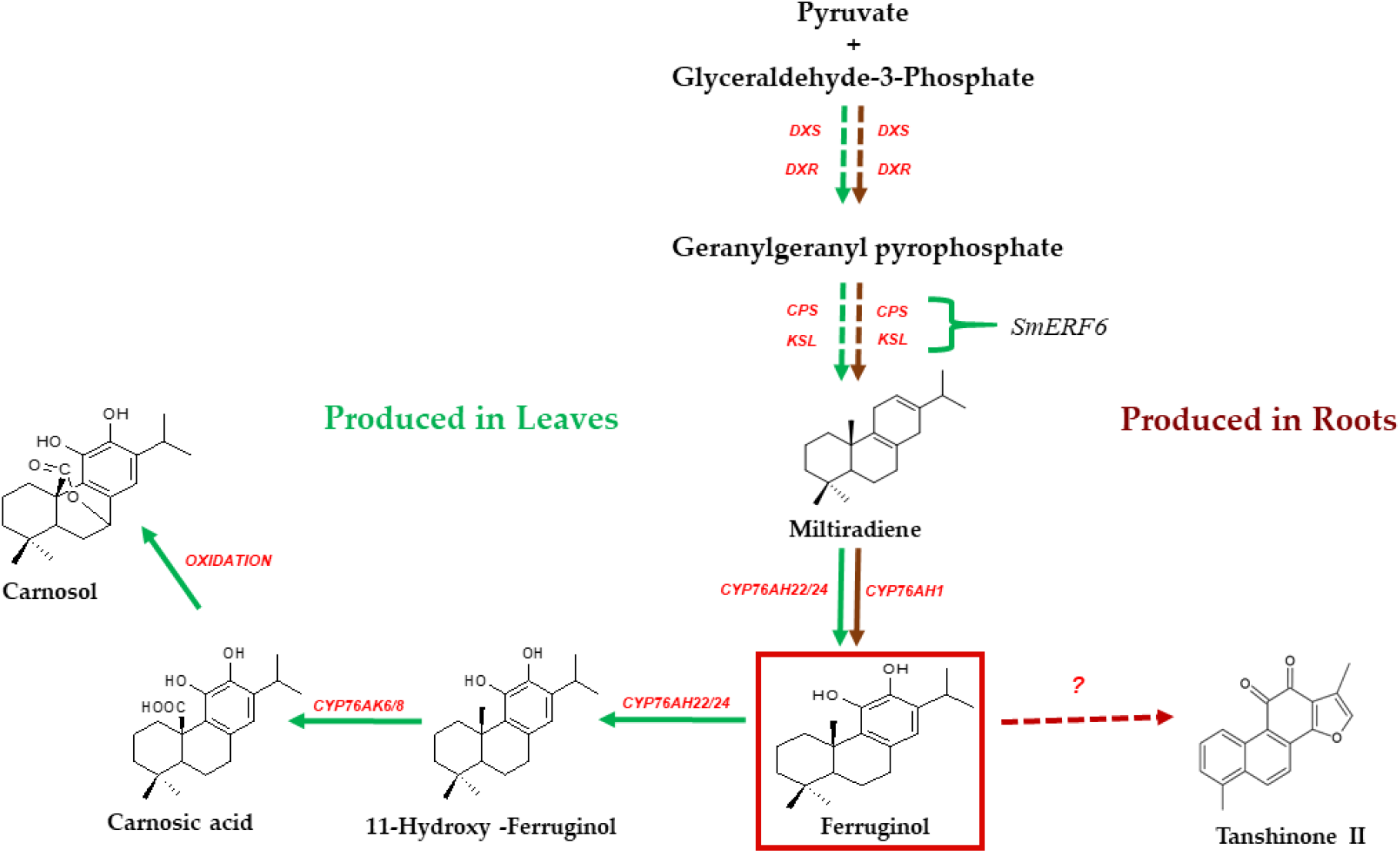
Schematic representation of diterpene diversity in *S. officinalis* and *S. miltiorrhiza*. CO and CA are produced in leaves of *S. officinalis* and tanshinones are produced in roots of *S. miltiorrhiza*. Ferruginol acts as a common precursor for the biosynthesis of CO, CA and tanshinones (Hu *et al*., 2021). *Sm*ERF6 is previously reported to bind on the promoter regions of *CPS* and *KSL* genes to increase their transcription level which in turn results in improved flux of ferruginol (Bai *et al*., 2018).

TFs, are generally known as trans-acting factors which bind to the cis-acting elements in the promoter region of a gene and regulate its expression (Goossens *et al*., 2017). ERF TFs are one of the largest class of TFs and their AP2 DNA binding domain is considered to be plant specific and does not occur in other higher eukaryotes (Feng *et al*., 2020). ERF TFs are known to regulate physiology and metabolism of the plants at various levels. They do play specific and active role in the regulation of an array of genes governing the operation of multi-enzyme pathways related to specialized metabolism (Zhou & Memelink, 2016). For instance, ORCA TFs belong to the ERF family and regulate alkaloid biosynthesis in *Catharanthus roseus*, *Ophiorrhiza sp* and tobacco. GAME (Glycoalkaloid metabolism) TFs of Solanaceae are found to be the typical examples of ERF TFs which are known to play precise role in sterol biosynthesis (Shoji and Yuan, 2021; Feng *et al*., 2020). These salient findings indicate the universal functionality of TFs towards orchestrating the specialized metabolism of plants in homologous and heterologous hosts. In the present study, *SmERF6* TF was heterologously expressed in *S. officinalis* in order to understand its function in CO biosynthesis. Accordingly, wet-lab experimental work as well as studies involving bioinformatics tools were focused on to develop stable transgenic lines by employing *in planta Agrobacterium* mediated genetic transformation. Results of the present study, involving transient as well as stable expression of *SmERF6* TFs in garden sage, confirmed the significant upregulation of CO biosynthetic cascade which resulted in improved bioaccumulation of CA and CO. Findings of the present study are found to be the first in relation to the improvement of high value diterpene content in *S. officinalis* by expressing pathway specific TFs. Also, this study validates the concept of transcription factor engineering to improve the *in planta* content of specialized metabolites in medicinal plants.

## Experimental

### Plant Material, tissue specific, leaf developmental and methyl jasmonate treatments

*S. officinalis* plants were germinated from seeds and maintained at the plant containment facility. For tissue specific experiments, leaf, stem and roots were harvested. Similarly, for leaf developmental regulation studies, 1^st^, 2^nd^, 3^rd^, 4^th^ and 5^th^ leaf developmental pairs were harvested from well-grown plant. Age related experiments were carried out by harvesting leaves from the garden sage plant at 30, 60, 90 and 120 days of plant age. 200µM of methyl jasmonate (MeJA, Sigma Aldrich, USA) was prepared by dissolving 4.65µl of MeJA in 100 µl of 100% ethanol and diluted to 200µM with sterile deionized water. Later, leaves of garden sage were drenched with MeJA solution and the plants were covered. Leaf samples were harvested at 0, 2, 6, 12, 24 and 48 h post treatment. Leaves sprayed with sterile deionized water were taken as control. All the harvested tissues were frozen in liquid nitrogen and stored at – 80^0^C for further analysis.

### Prediction of sub-cellular localization of CO biosynthetic pathway cascade

Sub-cellular localization of the CO biosynthetic pathway enzymes like Copalyl diphosphate synthase (CPS), Kaurene synthase like (KSL), Hydroxy Ferruginol Synthase (HFS), CYP76AK6/8 and Ethylene Responsive Factor 6 (ERF6) were analysed through bioinformatics software such as iPSORT, WoLF PSORT, Predotar and ChloroP according to Kumar *et al*. (2020).

### Generation of overexpression and silencing vectors

The coding sequences of *SmERF6* (Genbank accession number: KY988300.1; Bai *et al*., 2018) were de novo synthesized from Synbio Technologies, USA and cloned to pJET1.2/vector. The sequences were confirmed with sanger sequencing and sub-cloned to pBI121 binary vector in *Xba*I and *Sac*I sites. Later, the generated pBI121::*SmERF6* (overexpression), pBI12::*SmERF6AS* (silencing) constructs along with pBI121 (empty vector) were moved to *Agrobacterium tumefaciens* strain GV3101 by freeze thaw method. The primers used in the study were given in the Table S1.

### Transient expression of SmERF6

Transient expression studies were performed according to Dwivedi *et al*. (2022) with minor modifications. *A. tumefaciens* strains harbouring pBI121::*SmERF6*, pBI121::*SmERF6AS* and empty vector (pBI121) were grown overnight in Luria bertani (LB) broth (HiMedia, India) with respective antibiotics. Overnight grown cultures were pelleted down and re-suspended in MES [2-(N-morpholino) ethanesulfonic acid] buffer (50 mM MES pH 5.6, 2 mM Na_3_PO_4_, 0.5% glucose, 100 μM acetosyringone, Tween20) till the final OD_600_ reaches 0.2. Later, the *Agrobacteria* suspension was incubated for two hours at 28^0^C followed by infiltration. Garden sage twigs were excised from the 3-4 month old plants and leaves were scratched on the abaxial side with a needle less syringe. The twigs were placed upside down to submerge the leaves in *Agrobacteria* suspension and infiltrated under vacuum for 35 to 40 min in a desiccator (**Figure 5B**). Later, infiltrated sage twigs were kept in 5% sucrose solution and incubated at dark conditions for 48hrs. Post 48hrs, samples were collected and stored at – 80^0^C for further analysis.

### Generation of transgenic S. officinalis

*In planta Agrobacterium* mediated genetic transformation was performed according to Kumar *et al*. (2020). Overnight grown *A. tumefaciens* strain harbouring pBI121::*SmERF6* was pelleted and diluted in MES buffer until the OD_600_ reaches 0.2. Garden sage seeds were germinated in germination trays and cotyledonary leaf stage seedlings were employed for the experiments. Straight cut was made between both cotyledonary leaves to make a wound to expose apical meristematic tissue (**Figure 10A**). A sterile cotton swab was wrapped around the cotyledons to keep them intact and sealed with cellophane. *Agrobacteria* suspension of 20µL was added to the wounded site. The germination trays containing the plants were covered to maintain moisture and incubated at dark conditions. The *Agrobacteria* treatment was performed continuously for 4 days and on the 5^th^ day, old cotton swab was removed and new cotton was wrapped around the treated plants. Later, plants were treated with 300mg/L of cefotaxime for three consecutive days to remove the *Agrobacteria* contamination. After three days, the cotton was removed and the infected plants were maintained at plant containment facility. From one month old *Agrobacteria* infected plants (8-10 leaf stage), second pair of leaves were excised and subjected to genomic DNA isolation following the protocol of Tamari *et al*. (2013). Transgenic lines were confirmed with PCR analysis with *35S* forward and *SmERF6* reverse primers. To rule out the presence of *Agrobacterium* contamination, PCR analysis was performed with Chromosome virulence gene (*Chv*) primers. The primers used in the study were given in the Table S1.

### Total RNA isolation and gene expression analysis

Total RNA was isolated using Spectrum^TM^ Total Plant RNA isolation kit (Sigma Aldrich, USA) by following manufacturer’s protocol. First-strand cDNA was synthesized from total RNA using iScript^TM^ cDNA synthesis kit (BIO-RAD, USA) following the manufacturer’s instructions. Real-Time-quantitative-PCR (RT-qPCR) was performed using iTaq Universal SYBR Green supermix (BIO-RAD, USA) in the CFX Connect Real-time PCR system (BIO-RAD, USA). The primers used for this study were given in the Table S1. Each reaction was carried out in triplicates. RT-qPCR conditions were performed as follows: one cycle of 94^0^C for 10 min; followed by 40 cycles of 94^0^C for 15 secs and 60^0^C for 1 min. Fold change differences in gene expression were analyzed through comparative cycle threshold (*C_t_*) method (CFX Connect Real-time PCR detection system, BIO-RAD, USA). The *C_t_* for each gene was normalized with elongation factor A (*elfA*). Fold changes of gene expression were calculated using the relative quantification 2^-ΔΔCt^ method.

### Quantification of CO and CA

Metabolite extraction and quantification was performed according to Loussouarn *et al*. (2017). Leaf tissues were collected at same age and developmental stage for all the experiments. Empty vector infiltrated leaves were taken as controls for transient expression experiments and wild type plant leaves were taken as controls for stable transgenic experiments. Leaf tissue (100 mg) was grounded in 1ml of 99.5:0.5 v/v of methanol (HPLC grade, Merck, USA) and phosphoric acid (HPLC Grade, Sigma-Aldrich, USA). The extracts were centrifuged at 13,000 RPM for 15 min at 4^0^C and concentrated in a Rotavac evaporator (Concentrator Plus, Eppendorf, Germany). Later, concentrated extract was diluted in 100µL of HPLC grade methanol and filtered through a 0.45μm membrane filter (Merck, USA). The extracts were subjected to reverse phase HPLC and isocratic method was followed. Mobile phase for the analysis consisted; 65:34.8:0.2 of acetonitrile: water: phosphoric acid at a flow rate of 1ml/ min with a run time of 30 minutes. C18 reverse-phase column (Sunfire® 5 µm, 4.6 X 250 mm; Waters, Milford, MA, USA) was used for analysis in the HPLC system (Waters, USA). The compounds were detected through Ultra Violet (UV) wavelength at 230 nanometres (nm) and quantified using authentic standards of CO and CA (Sigma Aldrich, USA).

### Liquid Chromatography – Mass spectrometry Profiling

Leaf tissue (100 mg) was grounded in 1ml of methanol (LC-MS grade, Merck, USA). The extracts were centrifuged and concentrated in a Rotavac evaporator (Concentrator Plus, Eppendorf, Germany). Later, dried extract was diluted in 100µL of LC-MS grade methanol and filtered through a 0.45μm membrane filter (Merck, USA). LC-MS for the leaf extracts was performed using LC-MS/MS instrument (Waters, AcQuity H-Class Plus-UPLC instrument, USA) coupled with Xevo TQ-S cronos mass spectrometer (Waters, USA) with HESI (Heated electrospray interface) source. The column used was ACQUITY UPLC HSS T3 (2 × 100mm, 1.8 µm particle size) from Waters, USA, and oven temperature was maintained at 40°C. The mobile phase used was a gradient of water (A), and 90% acetonitrile (B); the gradient method was as follows: 0 min, 90% A, 10% B; 2 min, 90% A, 10% B; 5 min, 50% A, 50% B; 7 min, 10% A, 90% B; 9 min, 10% A, 90% B; 10 min, 90% A, 10% B; 12.50 min, 90% A, 10% B. The flow rate was maintained at 0.35 mL/min. The mass spectrometer was run in both negative and positive ion modes with a mass range of m/z 100–1000 at a resolution of 60,000 at m/z 100. The analyses were performed using MassLynx Mass Spectrometry Software (Waters, USA). The detection and identification of the compounds were analyzed by preparing a mass library of the compounds of interest and relative abundance was calculated by comparison with empty vector controls.

MS^2^ analysis for CO and CA standards along with plant extracts were performed according to the given protocol. The mass spectrometer was run in negative ion mode with a mass range of m/z 100–1000 at a resolution of 60,000 at m/z 100. For the MS^2^ measurements, collision-induced dissociation was kept at 20 eV. MS^2^ measurements were executed at m/z of CO (329.37) and CA (331.36) [M − H]− with a scan range of m/z 450–600 for both the standards and plant extracts. The analyses were performed using MassLynx Mass Spectrometry Software (Waters, USA).

### Gas Chromatography – Mass spectrometry Profiling

The volatiles in infiltrated leaves of *S. officinalis* were entrapped using SPME (solid phase micro-extraction) fibre. Infiltrated leaves were ground to powder using liquid nitrogen and the powder was transferred to glass vials immediately followed by sealing to avoid the escape of volatiles, SPME fibre was inserted in the glass vial and incubated at 37^0^C for 2 hrs. Later, SPME fibre was inserted into GC-MS for volatile detection. The metabolite analysis was carried out in a GC-MS instrument (Agilent Technologies GC 7890B and MS 5977B, USA) and helium was used as a carrier gas to separate the metabolites in an HP-5 MS column. The splitless mode was used for operation. The temperature of the injector was set to 280°C and the oven temperature was set at 80°C for 1.5 min, then ramped to 220°C at a rate of 10°C/min without holding, then increased to 310°C at a rate of 20°C/min held for 10 minutes and with a 5-minute solvent delay. The flow rate across the column was 1mL/min. The conditions for the operation of the mass spectrometer were set as follows: ion source temperature - 230°C, MS Quad temperature - 150°C, electron energy (70eV) and scanning range of m/z, 25-1000 amu. The identification of metabolites was done by matching the mass spectra of each compound from the library (3:1, signal: noise) using the NIST-17.L mass spectral library. For the mass spectra comparison, the matching value of the metabolite identity was set to more than 70. For co-elution, the mass spectra of all peaks were analyzed at three different points, the beginning, middle and end of each peak width. There was no co-elution seen in any of the identified peaks. Furthermore, the compounds were identified based on their retention index, mass spectrum, the calculation and comparison of the GC retention index of a series of alkanes (C8–C30).

### Statistical analysis

Three independent biological replicates were taken for each experiment in the current study. Data presented are the average mean value of independent biological, technical and experimental replicates. Statistical significance between the experimental samples was calculated using unpaired Student’s *t*-test.

## Results

### Spatio-temporal regulation of CO biosynthesis in S. officinalis

Previous reports suggest that CO and CA are produced predominantly in *S. officinalis*, *R. officinalis* and *S. fruticosa* of Lamiaceae. However, tissue specificity and developmental regulation of CO biosynthesis in *S. officinalis* is not reported yet. In order to understand the molecular and biochemical dynamics of CO biosynthesis in garden sage, expression of CO biosynthetic genes that include *CPS*, *KSL*, *CYP76AH22/24* (Hydroxy ferruginol synthase) and *CYP76AK6/8* were analysed in leaves, stem and root tissues by RT-qPCR. Transcript abundance of *CPS*, *KSL*, *CYP76AH22/24* and *CYP76AK6/8* were significant in leaves and negligible in stem and roots (**Figure 3A; (i)**). In order to further confirm, we performed metabolite analysis by HPLC to simultaneously detect CO and CA (**Figure 3A; (ii), (iii)**). It was observed that CO and CA accumulated at very high amounts in leaves and were not detected in other tissues (**Figure 3A; Figure S1**). CO and CA that were detected in plant samples were authenticated by subjecting to MS^2^ analysis, and determining the fragments of individual masses, CO (329.37; [M − H]−) and CA (331.36; [M − H]−). Identified compounds from sage leaf extracts were fragmented by MS^2^ and the fragmentation pattern was confirmed with authentic standards (**Figure S2**). MS^2^ analysis revealed that the daughter mass value was 285.37 m/z for CO. Individual molecular masses of 244.36 and 287.33 m/z were observed for CA in leaf extracts that corresponded to the fragmentation pattern of pure compounds (**Figure S2**).

**Figure 3:**
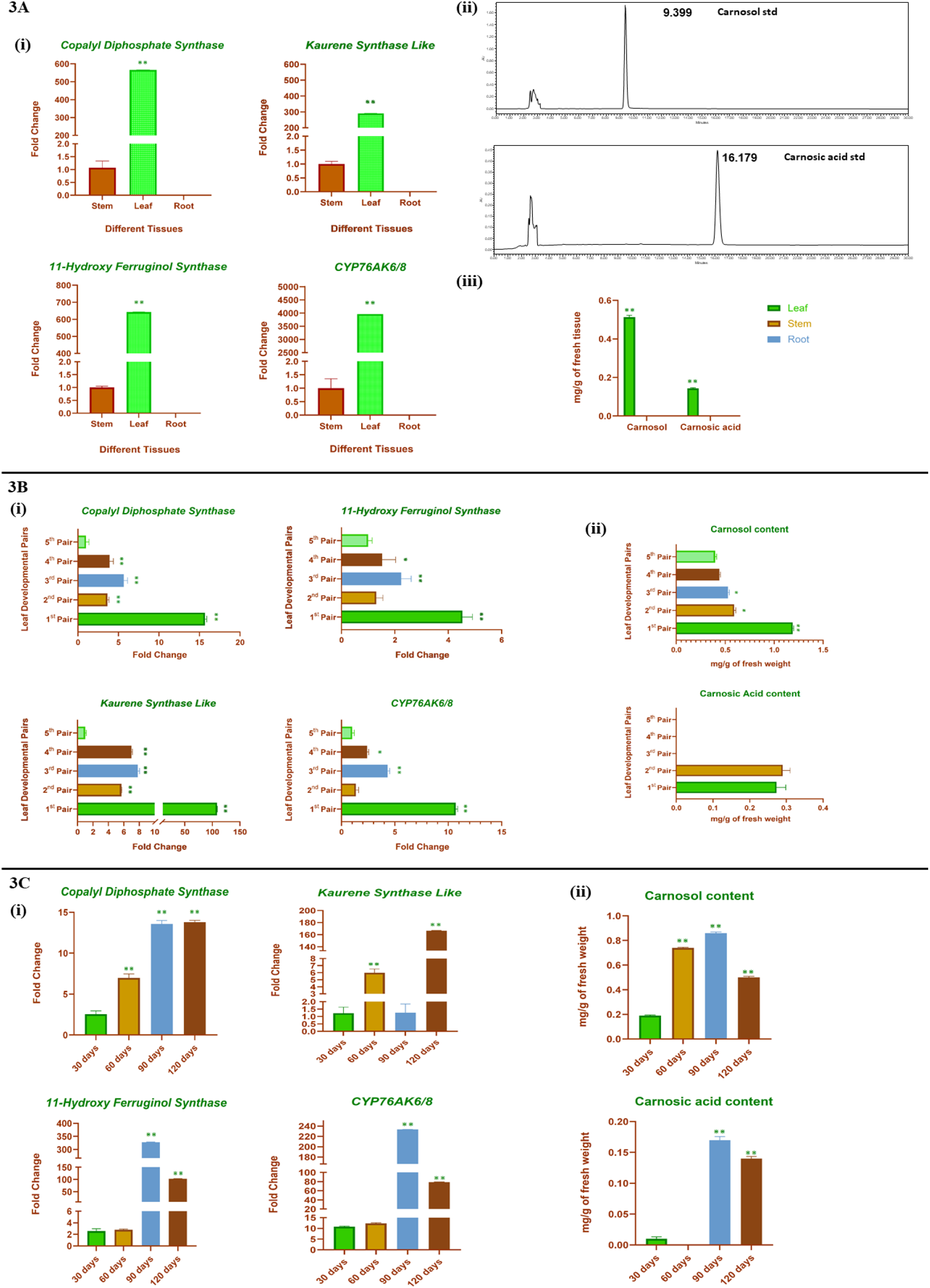
3A. Tissue specific expression of CO biosynthetic pathway genes and accumulation of CO and CA. (**i).** Expression analysis of CO biosynthetic pathway genes in leaf, stem and root tissues of *S. officinalis*. **(ii).** HPLC chromatograms of CO and CA standards. **(iii)**. Quantification of CO and CA accumulated in leaf, stem and root tissues of *S. officinalis*. **3B.** Leaf developmental pair regulation of CO biosynthesis in *S. officinalis*. **(i)**. Expression analysis of CO biosynthetic pathway genes in 1^st^ 2^nd^, 3^rd^, 4^th^ and 5^th^ leaf developmental pairs of *S. officinalis*. **(ii)**. Quantification of CO and CA accumulated in 1^st^ 2^nd^, 3^rd^, 4^th^ and 5^th^ leaf developmental pairs of *S. officinalis.* **3C.** Age dependent regulation of CO biosynthesis in *S. officinalis*. **(i)**. Expression analysis of CO biosynthetic pathway genes in leaves of *S. officinalis* post 30, 60, 90, 120 days. **(ii)**. Quantification of CO and CA accumulated in leaves of *S. officinalis* post 30, 60, 90, 120 days. Data presented are the average of three independent biological replicates. Significance (Student’s *t*-test) represented as \**p*< 0.05, \*\**p*< 0.01.

Age and developmental stage of plants are known to influence the metabolite profile. For example, young leaves accumulate higher amounts of metabolites as compared to their elder siblings in *S. fruticosa* and rosemary (Božić *et al*., 2015). In order to understand developmental stage specific regulation of CO biosynthesis in garden sage leaves, 1^st^, 2^nd^, 3^rd^, 4^th^ and 5^th^ developmental leaf pairs were subjected to gene expression analysis. In 1^st^ developmental leaf pair, significant transcript abundance of *CPS* (15-fold), *KSL* (100-fold), *HFS* (4-fold) and *CYP76AK6/8* (11-fold) genes were observed (**Figure 3B; (i)**). Gradual reduction in expression levels were observed in other developmental pairs (**Figure 3B; (i)**). Furthermore, metabolite analysis revealed abundant accumulation of CO and CA in 1^st^ developmental leaf pair as compared to other sets (**Figure 3B; Figure S3**).

Impact of plant age on the secondary metabolite profile was analysed by subjecting the 30-, 60-, 90-and 120-days old leaves to gene expression and metabolite accumulation analysis. Gene expression data revealed significantly higher levels of mRNA in 90 and 120-days old leaves when compared to early stages of the leaves (**Figure 3C; (i)**). Subsequently, HPLC analysis also confirmed the abundance of metabolites in 90 and 120-days old leaves (**Figure 3C; (ii)**). As a whole, these preliminary results indicated that leaves are the bio-factories of CO and CA in garden sage and metabolite accumulation is influenced by plant age and depends on the stage of leaf development. In addition, localization of CO biosynthetic cascade was predicted using different *in silico* tools, such as, Ipsort prediction, ChloroP, WOLFSPORT and Predotar. Maximum number of tools predicted the plastidial localization of CO biosynthetic cascade (**Table S2**).

### Methyl Jasmonate triggers the CO biosynthesis in S. officinalis

MeJA is a stress induced phytohormone that activates specialized metabolite producing biosynthetic cascades as a defence response. MeJA is conventionally used as an elicitor to trigger the specialized metabolite biosynthesis in plants. In the present study, we sprayed MeJA onto the garden sage leaves to understand its role in triggering CO biosynthesis. After treatment, *CPS* expression was induced post 2hrs (3 fold), 6hrs (13 fold) and 48hrs (4 fold). While, 5-fold and 3-fold enhanced expression of *KSL* was observed post 6 and 24hrs (**Figure 4A**). There was 5-fold and 4-fold improved transcript abundance of *HFS* which occurred post 6 and 24hrs of MeJA spray (**Figure 4A**). Expression of CYP76AK6/8 was found to be induced post 6hrs (4-fold) and 24hrs (3.5 folds) (**Figure 4A**). Observations showed that the CO biosynthetic cascade was induced upon MeJA treatment. Gene expression data was further validated by metabolite accumulation analysis, that revealed a 2-fold improved metabolite accumulation in post 6hrs MeJA treated garden sage leaves (**Figure 4B; Figure S4**).

**Figure 4:**
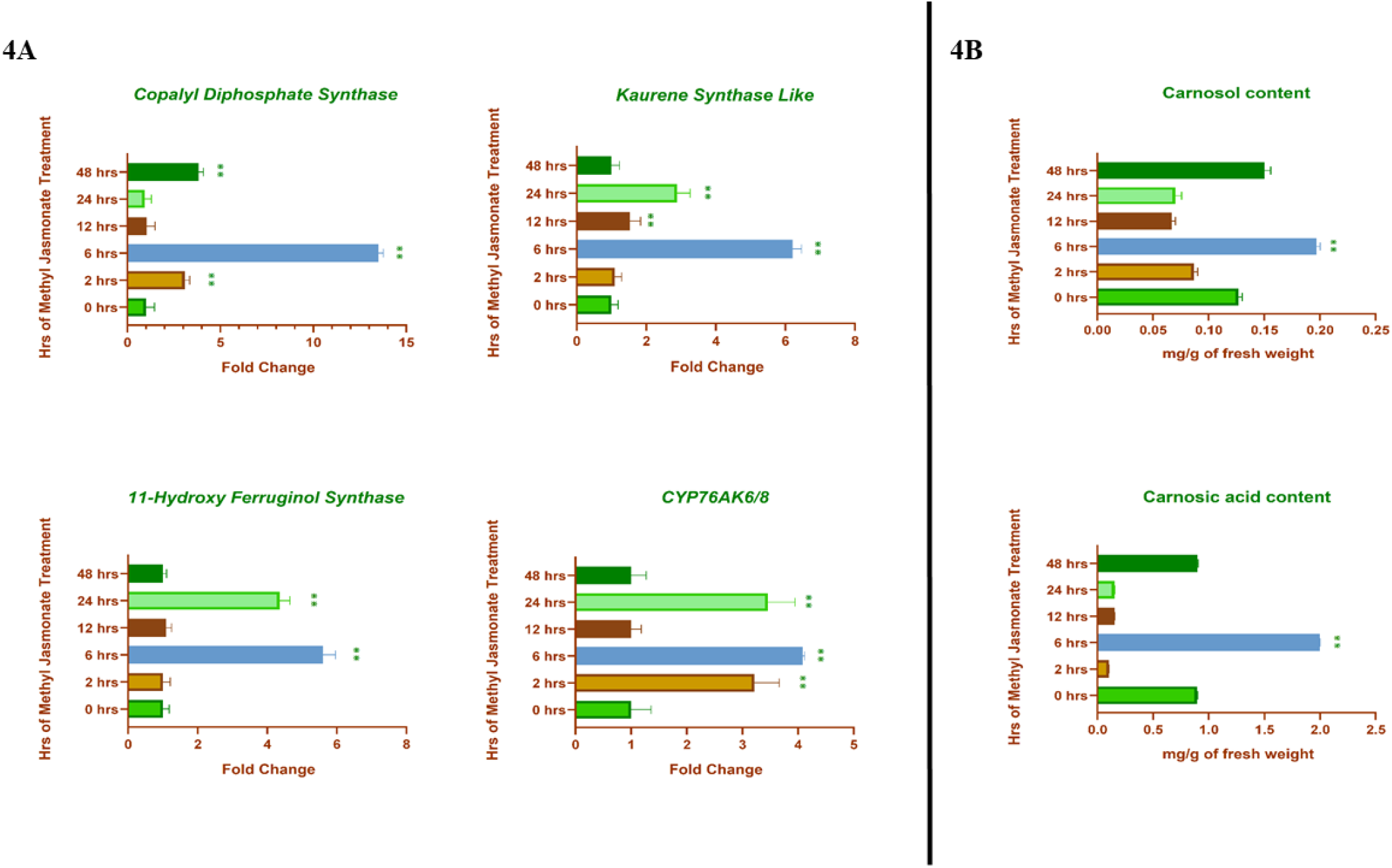
MeJA mediated induction of CO biosynthesis in *S. officinalis*. **4A**. Expression analysis of CO biosynthetic pathway genes at 0, 2, 6, 12, 24, 48 hrs post methyl jasmonate treatment in *S. officinalis*. **4B**. Quantification of CO and CA at 0, 2, 6, 12, 24, 48 hrs post MeJA treatment in *S. officinalis.* Significance (Student’s *t*-test) represented as \*\**p*< 0.01.

### Transient overexpression of SmERF6 induced CO biosynthesis in garden sage

Agroinfiltration is a rapid and robust technique that involves infusion of plant tissues with transformed *Agrobacteria* cells harbouring the gene of interest. This technique is being employed widely in order to produce the desired compounds in a short span of time and also to validate the gene function in homologous and heterologous host systems (Sathish *et al*., 2018). CA and CO are sparsely available in plant systems. In order to ensure abundant extraction possibilities for specialised metabolites from appropriate plant systems, manipulating the biosynthetic pathway by introducing an additional copy of pathway gene or transcriptional regulators to improve the metabolite production is found to be a reliable strategy. Here, we chose *SmERF6* as a potential candidate to improve metabolite accumulation, owing to its proven role in upregulating tanshinone biosynthesis (Bai *et al*., 2018). Accordingly, *Agrobacteria* cells harbouring overexpression (pBI121::*SmERF6*) and silencing constructs (pBI121::*SmERF6AS*) were infiltrated into garden sage leaves using vacuum (**Figure 5A** **& 5B**). Gene expression studies post 48hrs of infiltration displayed 5-fold increase in transcript abundance of *GGPPS* in infiltrated leaves as compared to empty vector controls (**Figure 6A**). mRNA expression of terpene synthases, *CPS* and *KSL* which are involved in conversion of GGPP to miltiradiene were improved to the extent of 10 and 12-folds, respectively (**Figure 6A**). Expression of two CYP450s, namely, *HFS* (11-fold) and *CYP76AK6/8* (9-fold), were enhanced in *SmERF6* infiltrated garden sage leaves (**Figure 6A**). In addition, salviol and sugiol which are the derivatives of ferruginol and are known to be synthesized through the catalytic activity of two CYP450s, namely, CYP71BE52 (salviol synthase) and CYP76AH1/3 (sugiol synthase) (Hu *et al*., 2021). In order to understand the diversion of ferruginol flux, we analysed the mRNA levels of salviol and sugiol synthases in infiltrated leaves, wherein it was observed that those two genes displayed increase of 2-fold and 6-fold, respectively (**Figure 6A**).

**Figure 5:**
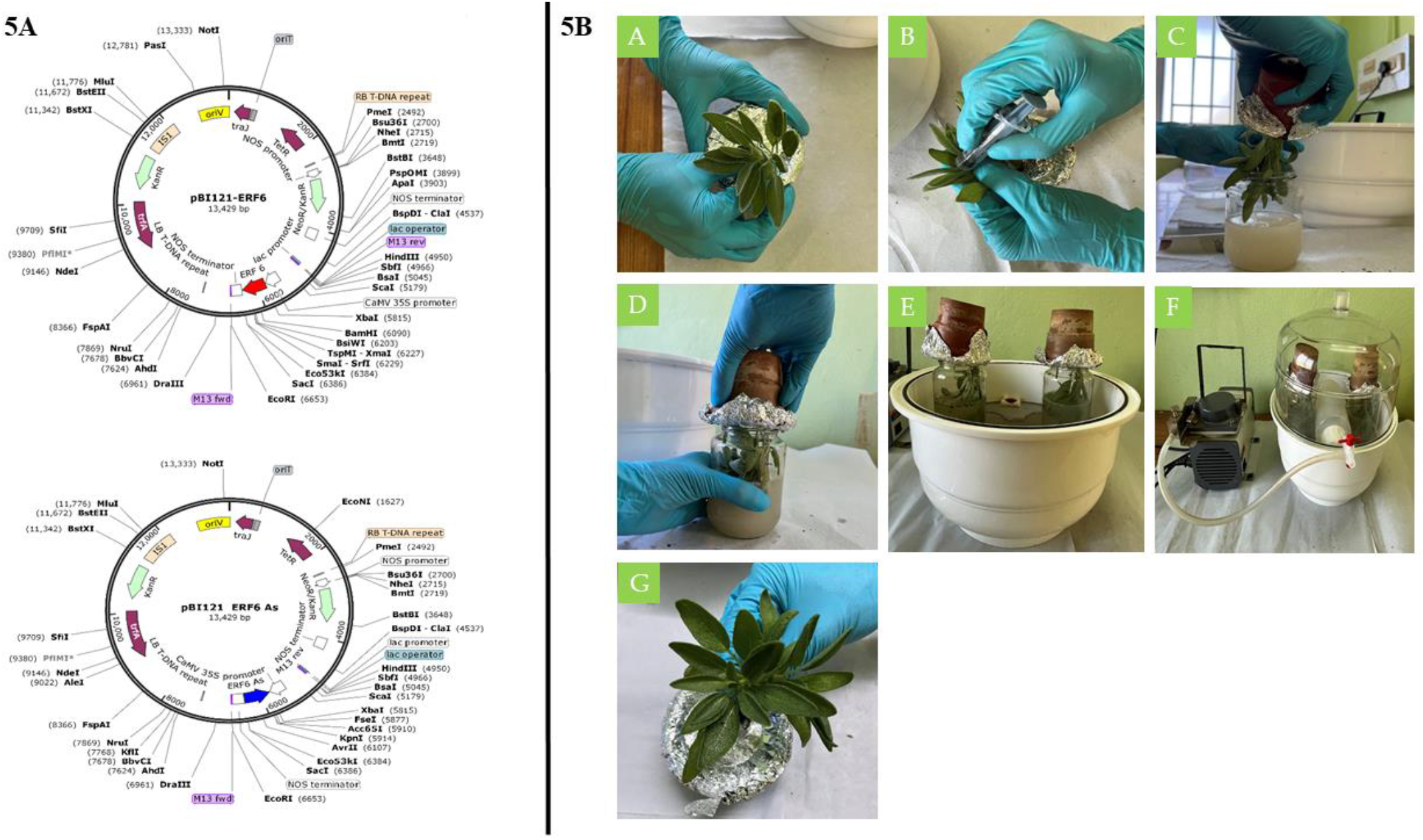
Vector maps of overexpression (pBI121::*SmERF6*), antisense (pBI121::*SmERF6AS*) constructs of *SmERF6* and vacuum infiltration procedure of *S. officinalis*. **5A**. Vector maps of overexpression (pBI121::*SmERF6*) and anti-sense constructs (pBI121::*SmERF6AS*) generated through SnapGene software (version: 3.2.1). **5B.** Schematic representation of vacuum infiltration procedure of *S. officinalis*. **A**: *S. officinalis* twigs were placed in a pot containing sterile soil and sealed with aluminum foil. **B**: Leaves were scratched with a needle less syringe on the abaxial side. **C, D**: Salvia twigs were immersed in *Agrobacterium* suspension harboring overexpression and antisense constructs separately. **E, F**: *S. officinalis* twigs were infiltrated using vacuum in a desiccator. **G:** *S. officinalis* leaves were removed from *Agrobacterium* suspension after infiltration and placed in 5% sucrose solution.

**Figure 6:**
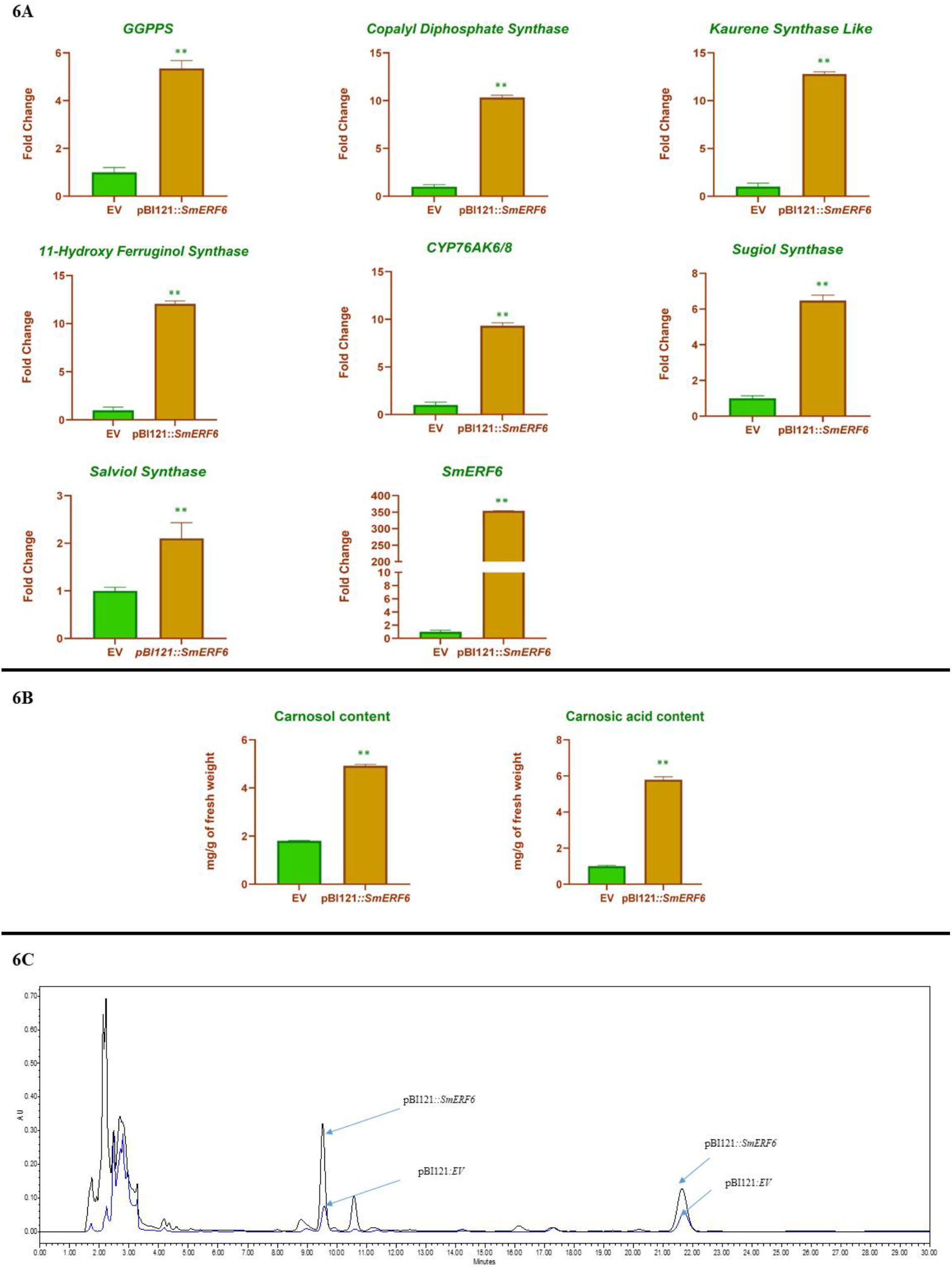
Expression analysis of CO biosynthetic pathway genes and HPLC analysis of CO and CA accumulation in pBI121::*SmERF6* infiltrated leaves of *S. officinalis*. **6A.** Relative expression of CO biosynthetic pathway genes in empty vector and *SmERF6* infiltrated *S. officinalis* leaves. *EFα* was used as internal reference gene for normalization. Relative gene expression levels of pBI121 control was set to 1. **6B**. Quantification of CO and CA in empty vector and *SmERF6* infiltrated *S. officinalis* leaves. Briefly, 100 mg of leaves were extracted thrice with methanol and phosphoric acid (99.5:0.5%) and subjected to HPLC. The peak area of CO and CA was normalized with internal standard. The amount of CO and CA in infiltrated leaves were calculated using authentic standards. **6C.** Representative HPLC chromatograms showing relative peaks of CO and CA in empty vector and *SmERF6* infiltrated leaves. Data presented were the average of three independent biological replicates. Significance (Student’s *t*-test) represented as \*\**p*< 0.01.

Significant increase in mRNA levels post infiltration were correlated with metabolite analysis through HPLC, wherein infiltrated leaves were enriched with 2-fold and 5-fold enhanced metabolite content, respectively (**Figure 6B** **& 6C**). In order to further validate the role of *SmERF6* in CO biosynthesis, knockdown assays were performed. In those experiments, contrasting results were observed in pBI121::*SmERF6AS* infiltrated leaves as compared to the overexpression data (**Figure 7**). Knockdown of homologous *SmERF6* did not alter the expression level of GGPPS and *CPS* genes. On the other hand, transcript levels of *KSL*, *HFS* and *CYP76AK6/8* genes were found to be drastically reduced (**Figure 7A**). In addition, it was observed that CA and CO levels were decreased in *SmERF6AS* infiltrated leaves (**Figure 7B**).

**Figure 7:**
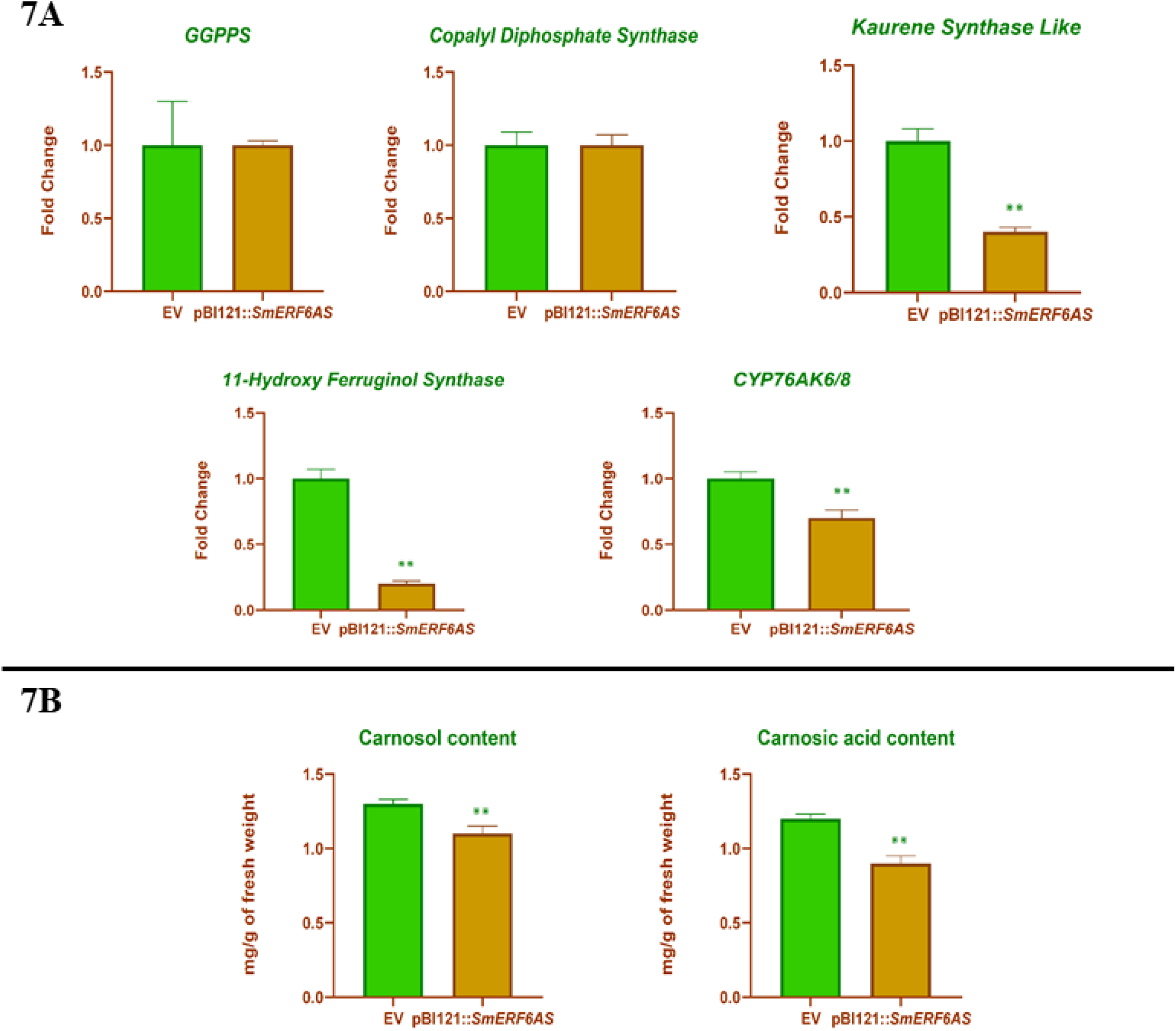
Expression analysis of CO biosynthetic pathway genes and HPLC analysis of CO and CA accumulation in pBI121::*SmERF6AS* (antisense constructs) infiltrated leaves of *S. officinalis*. **7A.** Relative expression levels of CO biosynthetic pathway genes in empty vector and *SmERF6AS* infiltrated *S. officinalis* leaves. *EFα* was used as internal reference gene for normalization. Relative gene expression levels of pBI121 control was set to 1. **7B**. Quantification of CO and CA in empty vector and *SmERF6AS* infiltrated *S. officinalis* leaves. Briefly, 100 mg of leaves were extracted thrice with methanol and phosphoric acid (99.5:0.5%) and subjected to HPLC. The peak area of CO and CA was normalized with internal standard. The amount of CO and CA in infiltrated leaves were calculated using authentic standards. The data given was the average of three independent biological replicates. Significance (Student’s *t*-test) represented as \*\**p*< 0.01.

LC-MS analysis was performed to understand the differential accumulation of other signature metabolites in garden sage subsequent to the introduction of *SmERF6*. A total of 15 various terpenes were detected from chromatogram along with other metabolites and the compounds were confirmed for their identity by respective m/z values and their corresponding mass spectrum (**Figure S5**; **Table S3**). Initially, significant increase in ferruginol content (62%) was observed in infiltrated leaves followed by enhanced accumulation of salviol (21%) and sugiol (600%) as compared to the empty vector controls (**Figure 8**). In garden sage, CA gets converted to its isoforms, such as, rosmadial, rosmanol, epirosmanol and methoxy carnosic acid through various enzyme mediated conversions (Birtić *et al*., 2015). Transient overexpression of *SmERF6* led to improved accumulation of CA derivatives. For example, enhancement of methoxy carnosic acid (170%), rosmanol (800%), rosmadial (285%) and epirosmanol (29%) was observed in the infiltrated leaves (**Figure 8**). In addition, terpene content was improved post infiltration, resulting in the abundance of linalool (120%), caryophyllene (170%), viridiflorol (500%), betulinic acid (95%) and asiatic acid (55%) when compared to appropriate controls (**Figure 8**). Later, we entrapped volatiles from infiltrated leaves and performed GC-MS analysis and a total of 32 compounds were detected (**Figure S6**; **Table S4**). However, relative abundance of volatiles did not alter much significantly post infiltration (**Figure 9**). These results draw a possible conclusion that ERF6 is highly specific to CO biosynthetic cascade and improves the content of CO and CA followed by its precursors and derivatives without affecting the biosynthesis of other terpenes which are essential for normal physiology of the plant.

**Figure 8:**
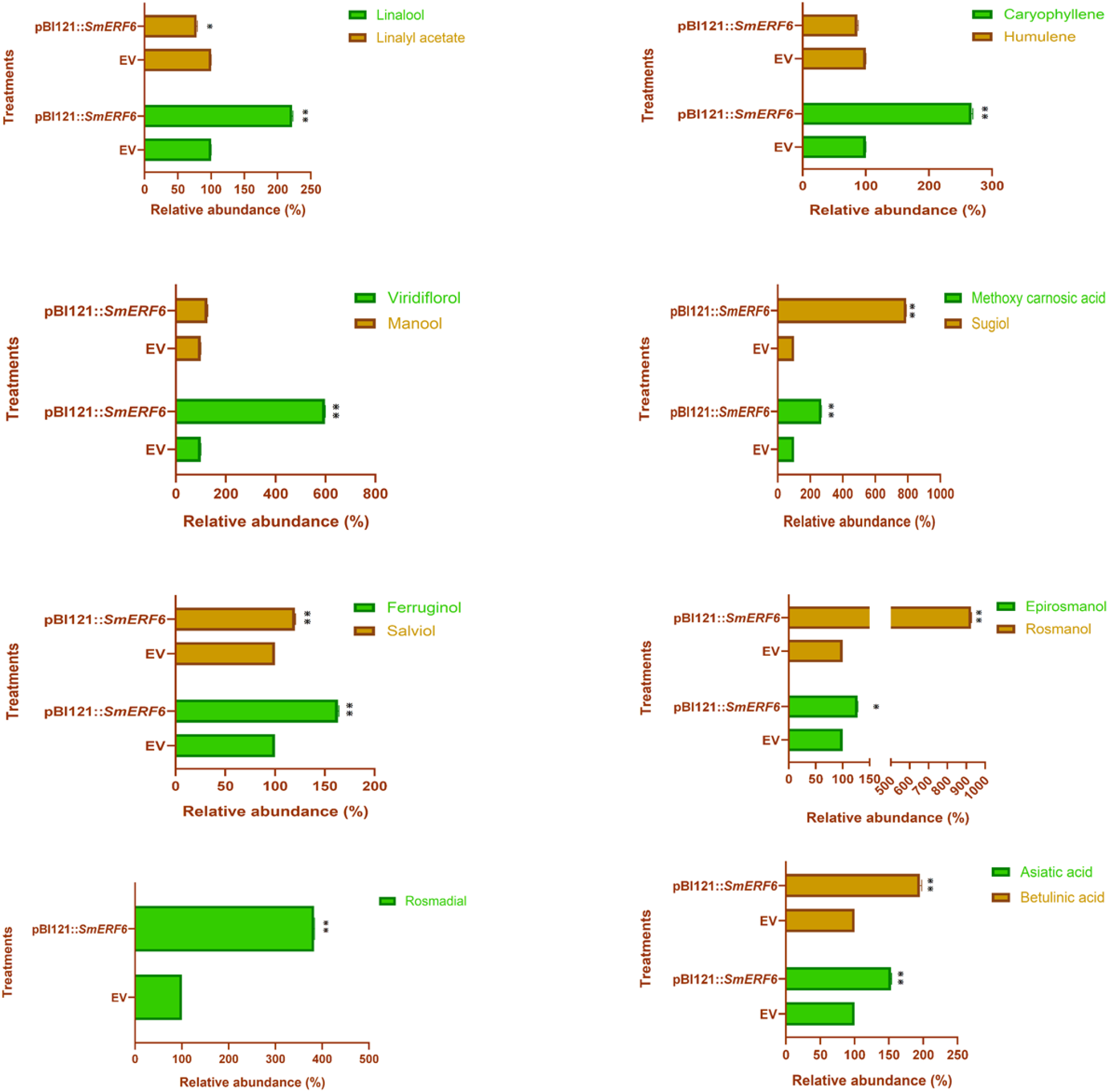
LC-MS profile of empty vector and *SmERF6* infiltrated *S. officinalis* leaves. Relative abundance of plant specific terpenes in *SmERF6* infiltrated leaves were calculated by comparing with empty vector data. Briefly, 100 mg of leaves were extracted thrice with methanol and subjected to LC-MS in both positive and negative ion modes. Data presented were the average of three independent biological replicates. Significance (Student’s *t*-test) represented as \*\**p*< 0.01. Monoterpenes: Linalool, linalyl acetate; sesquiterpenes: caryophyllene, humulene, viridiflorol; diterpenes: manool, methoxy carnosic acid, sugiol, salviol, ferruginol; CO derivatives: epirosmanol, rosmanol, rosmadial; triterpenes: Asiatic acid, betulinic acid.

**Figure 9:**
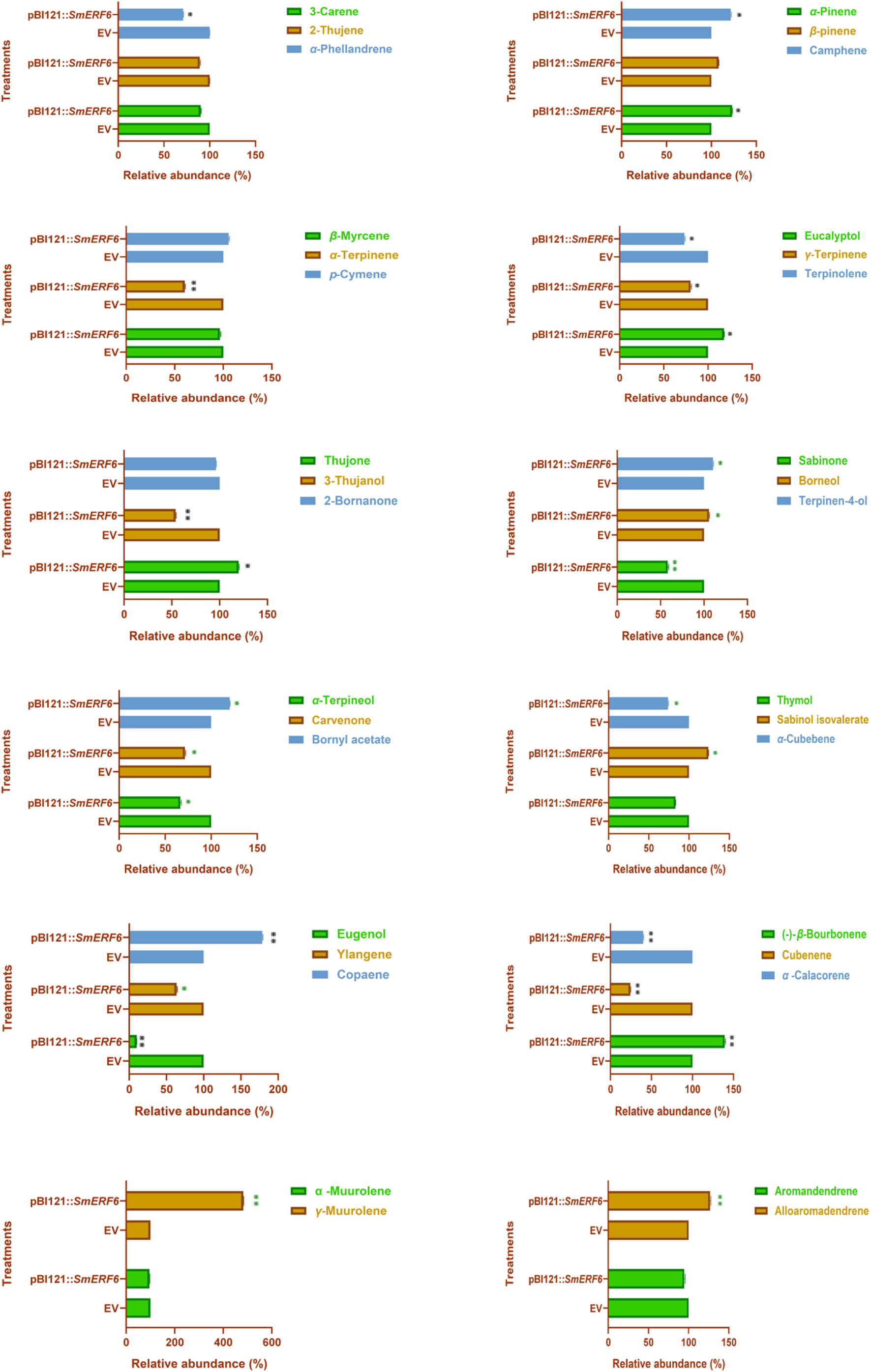
GC-MS profile of empty vector and *SmERF6* infiltrated *S. officinalis* leaves. Relative abundance of volatile terpenes in *SmERF6* infiltrated leaves were calculated by comparing with empty vector data. Volatiles from the *SmERF6* and empty vector infiltrated leaves were entrapped by SPME fiber and analyzed through GC – MS. Compounds were identified based on their retention index, comparing the mass spectrum of the components to the mass spectral library from NIST 17. L (National Institute of Standards and Technology, USA). Data presented were the average of three independent biological replicates. Significance (Student’s *t*-test) represented as \**p*< 0.05, \*\**p*< 0.01.

### Transgenic overexpression of SmERF6 improved CO accumulation in garden sage

Transient expression analysis results were further authenticated by generating stable transgenics of *S. officinalis* overexpressing *SmERF6*. Since, *S. officinalis* is a hardy plant and recalcitrant to *in-vitro* tissue culture techniques, a tissue culture independent *in planta Agrobacterium* mediated genetic transformation method was employed to generate transgenic *S. officinalis* (**Figure 10A**). Subsequent to subjecting to experimental conditions, genomic DNA from well grown putative transgenic plants were isolated and subjected to PCR analysis using *35s* forward and *SmERF6* reverse primers in addition to *Chv* primers to check *Agrobacterium* contamination (**Figure S7**). Out of 347 *Agrobacterium* infected plants, 115 plants survived and were screened for the functional integration of the transgene. Post PCR analysis, only three plants were found to be positive for transgenic nature (**Figure 10B**). The transformation efficiency of the experiment was derived to be 4.3% (**Table S5**). Initially, the expression of *SmERF6* was analysed in transgenic lines and it was observed that TR2 line (28-fold) displayed highest expression followed by TR3 (22-fold) and TR1 (18-fold) lines (**Figure 11A**). Hence, TR2 line is considered as highest expressing line, TR3 as moderate line and TR1 as least expressing transgenic line. Gene expression studies revealed similar pattern of expression levels as compared to the transient expression analysis experiments, wherein 2-3 folds improved mRNA levels of GGPPS were observed in transgenic lines (**Figure 11A**).

**Figure 10:**
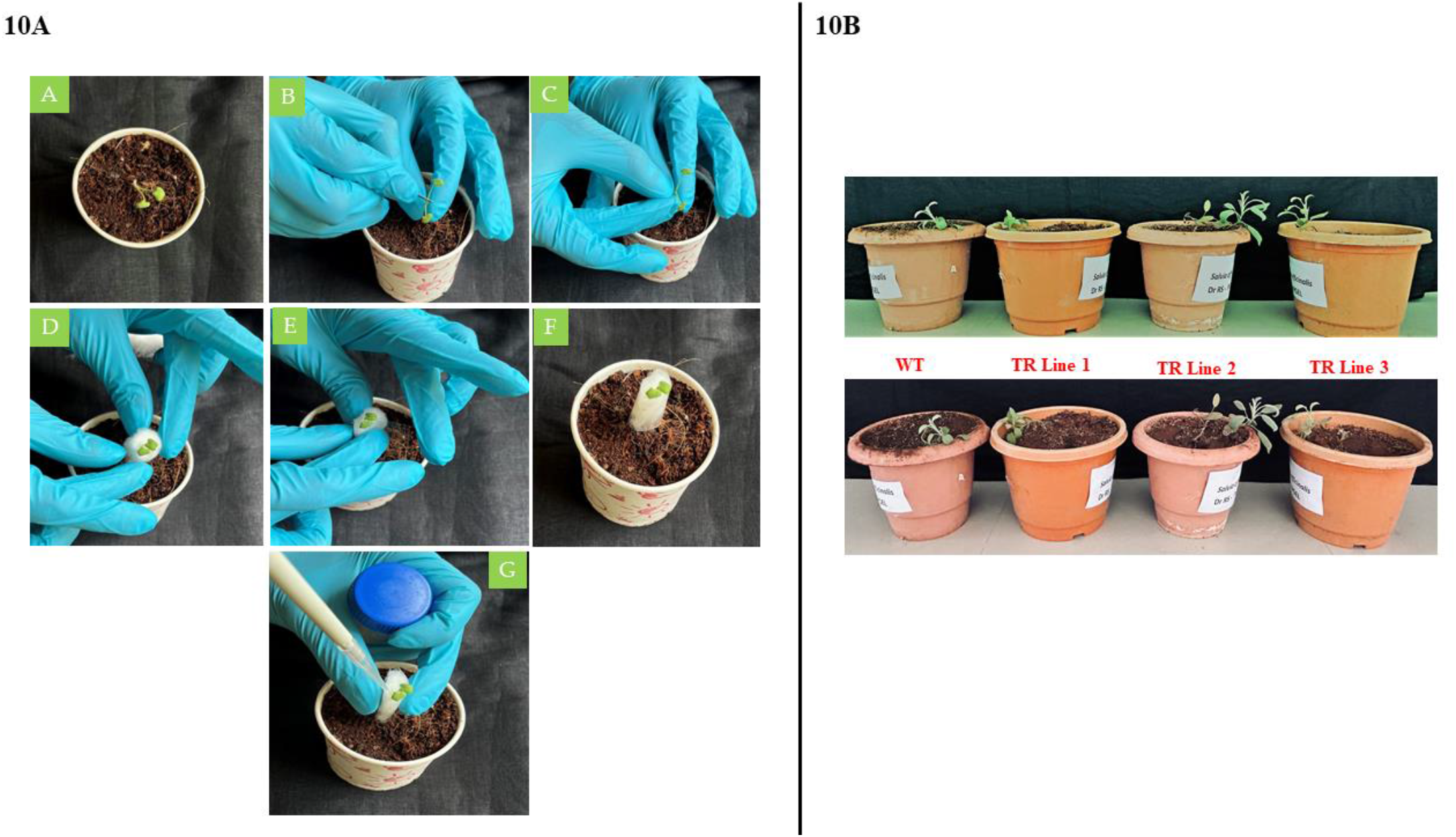
Schematic representation of *in planta Agrobacterium* mediated genetic transformation method. **10A. A.** Cotyledonary stage of *S. officinalis*; **B, C.** Wound was made in between the cotyledons using a sterile scalpel blade. **D, E, F.** Cotyledons were kept intact by using sterile cotton and wrapped with tape. **G.** *Agrobacterium* cultures harboring pBI121::*SmERF6* was added at the wounded site. **10B.** Photographic images of wild type and transgenic *S. officinalis* expressing *SmERF6* transcription factor.

**Figure 11:**
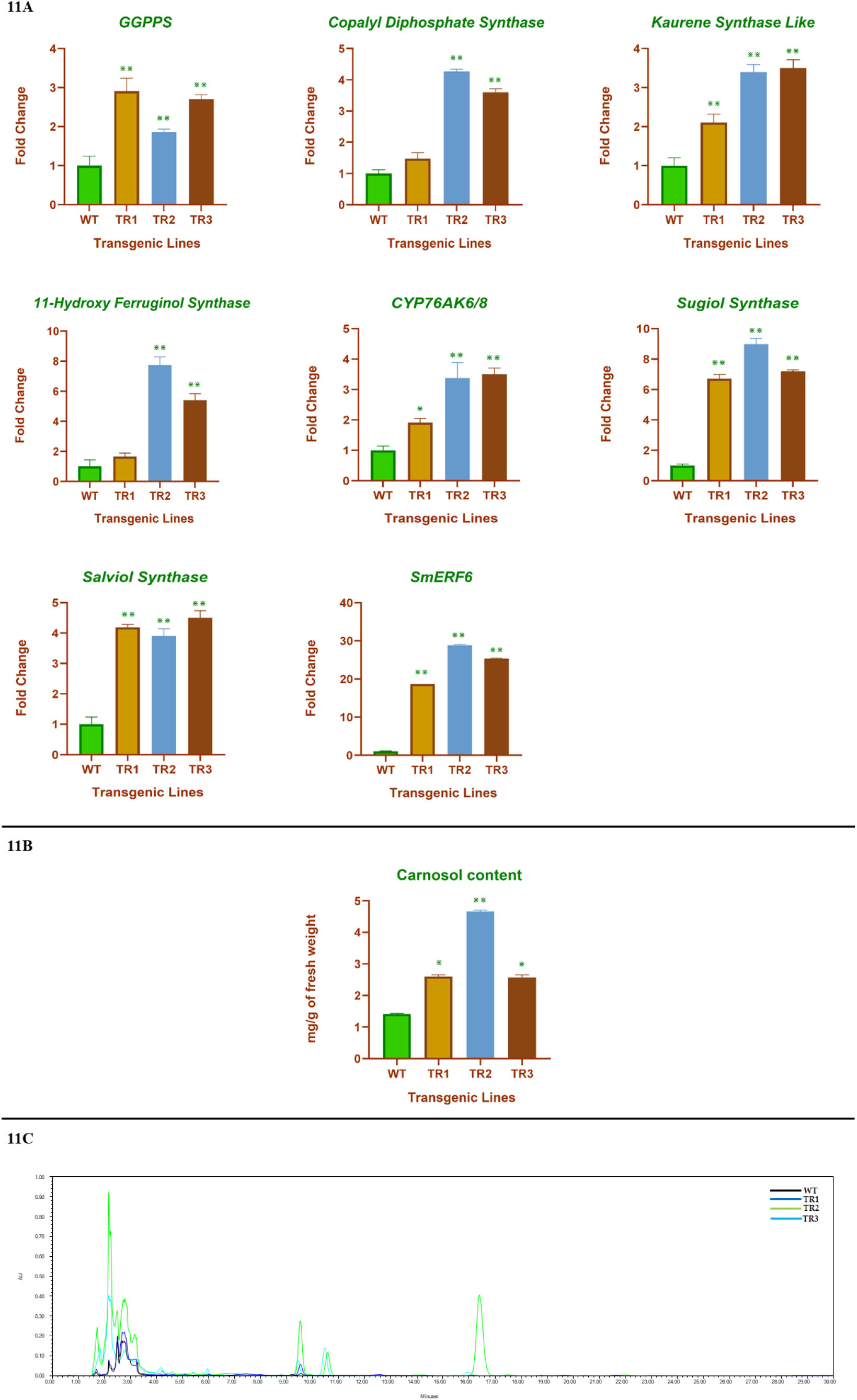
Expression analysis of CO biosynthetic pathway genes and HPLC analysis of CO and CA accumulation in wild type and transgenic *S. officinalis* expressing *SmERF6*. **11A.** Relative expression of CO biosynthetic pathway in wild type and transgenic *S. officinalis* expressing *SmERF6*. *EFα* was used as internal reference gene for normalization. Relative gene expression levels of wild type were set to 1. **11B**. Quantification of CO and CA in wild type and transgenic *S. officinalis* expressing *SmERF6*. Briefly, 100 mg of leaves were extracted thrice with methanol and phosphoric acid (99.5:0.5%) and subjected to HPLC. Peak area of CO and CA was normalized with internal standard. Amount of CO and CA in transgenic *S. officinalis* lines were calculated using authentic standards. **7C.** Representative HPLC chromatograms showing relative peaks of CO and CA in wild type and transgenic *S. officinalis* expressing *SmERF6*. Data presented were the average of three independent biological replicates. Significance (Student’s *t*-test) represented as \*\**p*< 0.01.

Further, 4-folds increased expression of CPS was observed in TR2 line followed by TR3 (3.5 fold) and TR1 (1.5 fold) lines (**Figure 11A**). It was also observed that 3.5-folds levels of transcript abundance of *KSL* gene occurred in TR2 and TR3 lines. While highest expression level of *HFS* was seen in TR2 (7.2 fold), moderate expression was observed in TR3 (5-fold) and least expression in TR1 (1.5 fold). Transcript abundance of *CYP76AK6/8* ranged between 2 to 4 folds in transgenic lines (**Figure 11A**). In addition, expression levels ranged between 6-8 folds and 4 to 4.5 folds for *Sugiol synthase* and *Salviol synthase,* respectively (**Figure 11A**). The CO accumulation trend in transgenic lines correlated with transcript data wherein 4-folds higher accumulation was observed in TR2 line followed by 2-folds in TR1 and TR3 (**Figure 11B** **& 11C**).

## Discussion

CO and CA are two high value diterpenes synthesized by the members of *Salvia* genus, *S. officinalis, S. fruticosa* and *R. officinalis*. It has been reported that biosynthesis of CO, CA is localized in specialized glandular trichomes on younger parts of the plant (Božić *et al*., 2015). In the present study, we observed the highest expression level of CO biosynthetic machinery and metabolite accumulation in garden sage leaves, which showed no detectable level of expression in stem and roots (**Figure 3A**). Our results are in concordance with previous reports on *S. fruticosa* and *R. officinalis* wherein the metabolites of the biosynthetic pathway were detected in leaves. Furthermore, in *S. fruticosa* and rosemary systems, gene expression was found to be minimal in the leaves that lacked trichomes, indicating the possible localization of CO biosynthesis (Brückner *et al*., 2014; Božić *et al*., 2015). In addition, *in planta* metabolite content correlated with the plant age and it was observed that expression of the metabolic cascades related to specialized metabolites seized upon aging of the plants in Lamiaceae members, such as, peppermint, *Ocimum sanctum* and rosemary (McConkey *et al*., 2000; Renu *et al*., 2014; Brückner *et al*., 2014). Comparably, our results indicated that CO biosynthesis is predominant in younger leaves of *S. officinalis* as compared to older leaves (**Figure 3B**). It is quite interesting to note that similar pattern of metabolite biosynthesis was observed in a related species, namely, *S. fruticosa* (Božić *et al*., 2015). In general, specialized metabolite biosynthetic cascades in higher plants get triggered upon MeJA treatment as a defense strategy. Further, previous reports confirm the increase of CA in *Salvia sclarea* and rosemary post MeJA treatment (Vaccaro *et al*., 2016; Yao *et al*., 2022). However, molecular mechanism beneath induction of genetic cascades related to CO biosynthesis upon MeJA spraying is not well understood in *S. officinalis*. In our experiments, it was observed that CO biosynthetic pathway was induced in *S. officinalis* post 6hrs of MeJA treatment (**Figure 4A, 4B**). Comparable to the observations of the present study, it has been shown that induction of tanshinone biosynthesis occurred post 12hrs of MeJA treatment in *S. miltiorrhiza* (Gao *et al*., 2014; Ge *et al*., 2015). Further, at natural conditions CO content ranges between 0.5 to 1 mg /gm fresh weight (approx. 0.001%) in leaves of *S. officinalis* (**Figure 3A**). Low level *in planta* availability of diterpenes in garden sage is inferred to be a major hindrance to extract the desired metabolites at commercial scale. Accordingly, we designed our work to improve the *in planta* availability of CO and CA by transcription factor engineering. Nevertheless, it can be noted that in-depth knowledge about the transcriptional regulators of CO biosynthesis across accumulators is currently unavailable.

Henceforth, we performed a comprehensive literature survey in order to obtain information related to the transcriptional regulators that are involved in tanshinone biosynthesis of *S. miltiorrhiza*. Tanshinones are abietane diterpenes, structurally similar to CO and derivatives of ferruginol (**Figure 2**). In total, we identified 45 transcription factors of different classes that actively participate in regulating the transcription of tanshinone biosynthetic cascades (unpublished data). In the current study, we chose *Sm*ERF6 that binds to the promoters of *CPS* and *KSL* genes in *S. miltiorrhiza* thus improving the flux towards the biogenesis of ferruginol, resulting in the accumulation of higher levels of tanshinones (Bai *et al*., 2018). In an earlier study, Bai *et al* (2018) observed overexpression of *SmERF6* in transgenic hairy roots of *S. miltiorrhiza* which led to improved transcript abundance of *CPS*, *KSL* genes (4 to 6-folds) along with a 3-fold augmentation of tanshinone content (Bai *et al*., 2018). Primarily, in our studies we transiently overexpressed *SmERF6* in garden sage through vacuum that resulted in 8 to 10-folds increase in expression of CO biosynthetic cascade (*GGPPS*, *CPS*, *KSL*, *CYP76AH22/24* and *CYP76AK6/8*) and 4-fold improved accumulation of CO and CA (**Figure 6A** **& 6B**). These results suggest that upstream pathway genes such as *GGPPS*, *CPS* and *KSL* could be the possible targets of *Sm*ERF6 and led to diverting the flux of precursors towards production of ferruginol. Observations of the present study indicated improvement of ferruginol content (60%) in the infiltrated leaves (**Figure 8**). Apart from CO, ferruginol acts as precursor for the production of sugiol and salviol in garden sage and other related species (Trikka *et al*., 2015; Ignea *et al*., 2016; Ali *et al*., 2017; Bajpai *et al*., 2021). Interestingly, mRNA levels of sugiol and salviol synthases were upregulated post infiltration along with improved metabolite levels (**Figure 6A**, **Figure 8**). Our results provide evidence that overexpression of *SmERF6* in garden sage improves the transcript abundance of *CPS* and *KSL* genes leading to significant levels of ferruginol. In addition, the results indicate that the accumulated metabolite pool is diverted towards the biosynthesis of CA, CO, salviol and sugiol by the catalytic activity of respective CYP450s. Additionally, flux of CA and CO were diverted to the production of respective derivatives, such as, methoxy carnosic acid, rosmanol, epirosmanol and rosmadial in infiltrated leaves (**Figure 8**). However, it is pertinent to mention that *SmERF6* infiltration did not alter the levels of volatile terpenes in garden sage (**Figure 9**). Moreover, knockdown of homologous *ERF6* drastically reduced the expression of CO biosynthetic genes and metabolite levels, thereby confirming the positive role of *ERF6* in metabolite production (**Figure 7A** **& 7B**). In order to substantiate transient experiment results, stable transgenic lines of *S. officinalis* overexpressing *SmERF6* were developed (**Figure 10**). The results obtained through gene expression and metabolite analysis of stable lines were consistent with transient experiments and displayed improved transcript abundance and metabolite content in stable transgenic lines as compared to the wild types (**Figure 11**).

## Conclusion

Transcription factors are the potential metabolic engineering tools to improve the *in planta* content of high value metabolites in various medicinal plants. In the present study, heterologous overexpression of *SmERF6* transcription factor in *S. officinalis* was engineered in order to understand its role in modulating CO biosynthesis. In garden sage, *SmERF6* improved transcript abundance of CO biosynthetic cascade that resulted in enhanced accumulation of CO and CA. In addition, comprehensive characterization of target genes of *SmERF6* in *S. officinalis* needs to be identified yet. To the best of our knowledge, this is the first report on improving high value diterpene content in *S. officinalis* by overexpressing pathway specific transcription factors. Also, we developed stable transgenics of *S. officinalis* overexpressing *SmERF6* by *in planta Agrobacterium* mediated genetic transformation which can be used to extract the cherishable metabolites at commercial scales in order to meet global market demands for the pharmaceutically important specialized metabolites.

## Author Contributions

RB performed the experiments, data analysis and prepared manuscript. GT assisted RB in performing cloning and RT-qPCR experiments. PD assisted RB in performing HPLC and GC-MS analysis. DS supervised PD. RS conceived the project and supervised RB, GT.

## Acknowledgments

RS acknowledge University Grants Commission, Government of India for financial support through UGC-BSR-Mid Career Grant (Sanction no. 19-266/2021(BSR)). RB acknowledge Dr. S. Rajeev Kumar, Scientist and Head, NPI division, String Bio Pvt Ltd, Bengaluru, India for technical help to carry out the work.

## Supplementary Figures

**Figure S1:** HPLC chromatograms of A. Leaf tissue; B. Stem tissue; C. Root tissue.

**Figure S2:** Chromatograms and mass fragmentation patterns of CO and CA through LC-MS analysis. HPLC chromatograms of **A.** CO standard (9.5 min); **B**. CA standard (16.2 min); **C**. Mass fragmentation of CO standard (m/z: 329.37); **D**. Mass fragmentation of CO in leaf extracts of *S. officinalis* (m/z: 329.37). **E**. Mass fragmentation of CA standard (m/z: 331.28); **F**. Mass fragmentation of CA in leaf extracts of *S. officinalis* (m/z: 331.28).

**Figure S3:** HPLC Chromatograms of different developmental stage leaf pairs extracts **A**. 1^st^ pair; **B**. 2^nd^ pair; **C**. 3^rd^ pair, **D**. 4^th^ pair; **E**. 5^th^ pair.

**Figure S4.** HPLC Chromatograms of sage leaf extracts post MeJA treatment. **A**. 0hrs, **B.** 2hrs, **C.** 6hrs, **D.** 12hrs, **E.** 24hrs, **F.** 48hrs.

**Figure S5.** Total ion chromatograms (TIC) of *S. officinalis* leaf extracts infiltrated separately with *A. tumefaciens* harbouring pBI121:EV and pBI121*::SmERF6* constructs. LC-MS was operated in both positive and negative ion modes to analyze the metabolite contents. **A**. Negative ion mode of pBI121:EV; **B**. Positive ion mode of pBI121:EV, **C.** Negative ion mode of pBI121*::SmERF6*, **D**. Positive ion mode of pBI121*::SmERF6*. The retention times of each detected metabolite and respective m/z values were provided in Table S3.

**Figure S6.** The GC-MS chromatograms of *S. officinalis* extracts infiltrated separately with *A. tumefaciens* harbouring pBI121:EV and pBI121::*SmERF6* constructs. A. pBI121:EV, B. pBI121::*SmERF6*. Retention times of each detected metabolite and respective m/z values were provided in Table S4.

**Figure S7.** Screening of putative transgenic lines of S. *officinalis* transformed with *SmERF6*. A. lane shows the PCR amplification of integrated gene with *35S* forward and *SmERF6* reverse primers; 01: wild type, 02-06: putative transgenic lines, 07: positive control, 08: GeneRuler Ladder mix (Thermo scientific, USA). B. Lane shows the PCR analysis to check the *Agrobacterium* contamination in putative transgenic lines by using *Chv* gene specific primers; 01: wild type, 02-06: putative transgenic lines, 07: positive control, 08: GeneRuler Ladder mix (Thermo Scientific, USA).

## Supplementary Tables

**Table S1:** Primers used in the study

**Table S2:** Sub-cellular localization of CO biosynthetic genes using various bioinformatics tools.

**Table S3:** List of compounds detected in leaf extracts of *S. officinalis* infiltrated with pBI121:EV and pBI121::*SmERF6* analyzed through LC-MS. The area of the compounds detected in respective samples were given in the table. Samples were run in triplicates and the data presented are average of three replicates.

**Table S4:** List of VOCs detected in leaf extracts of *S. officinalis* infiltrated with pBI121:EV and pBI121::*SmERF6* analyzed through GC-MS. The samples were run in triplicates and the data presented are average of three replicates.

**Table S5:** Efficiency of *in planta Agrobacterium* mediated genetic transformation in *S. officinalis*.

