## Supplementary information for "SmERF6 promotes the expression of terpenoid pathway in Salvia officinalis and improves the production of high value abietane diterpenes, carnosol and carnosic acid"

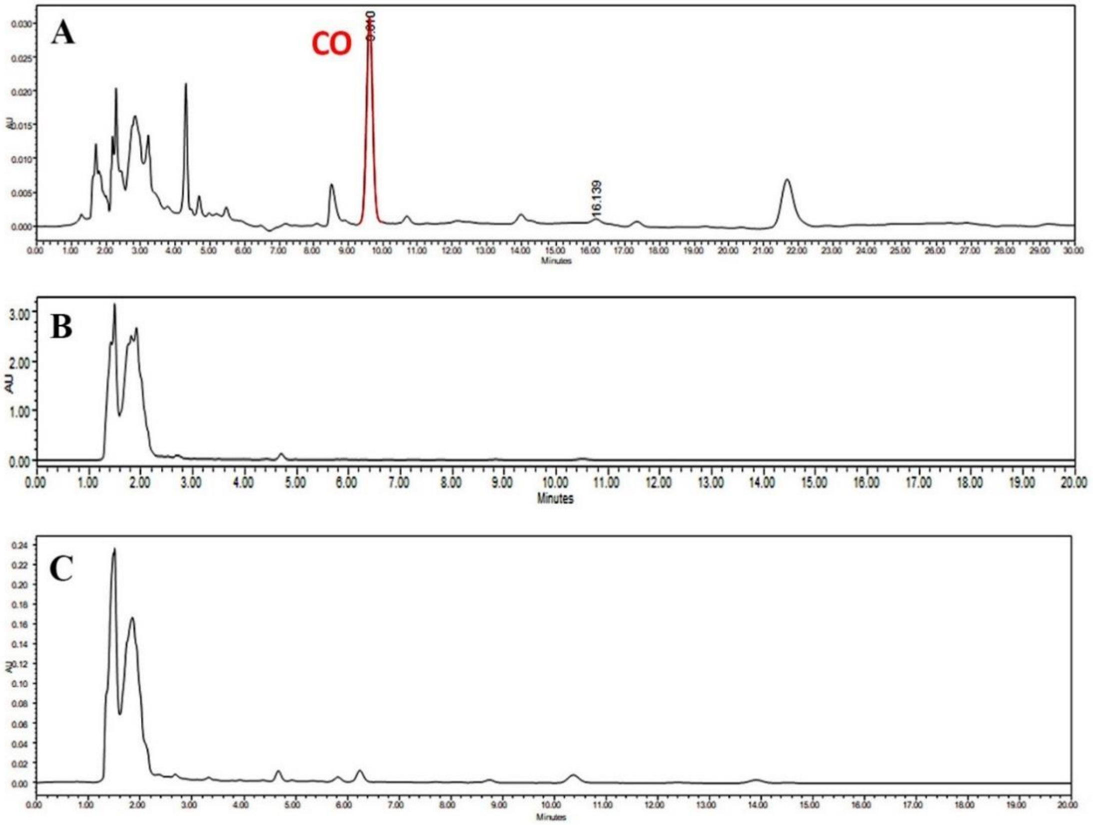


**Figure S1:** HPLC chromatograms of A. Leaf tissue; B. Stem tissue; C. Root tissue**.**

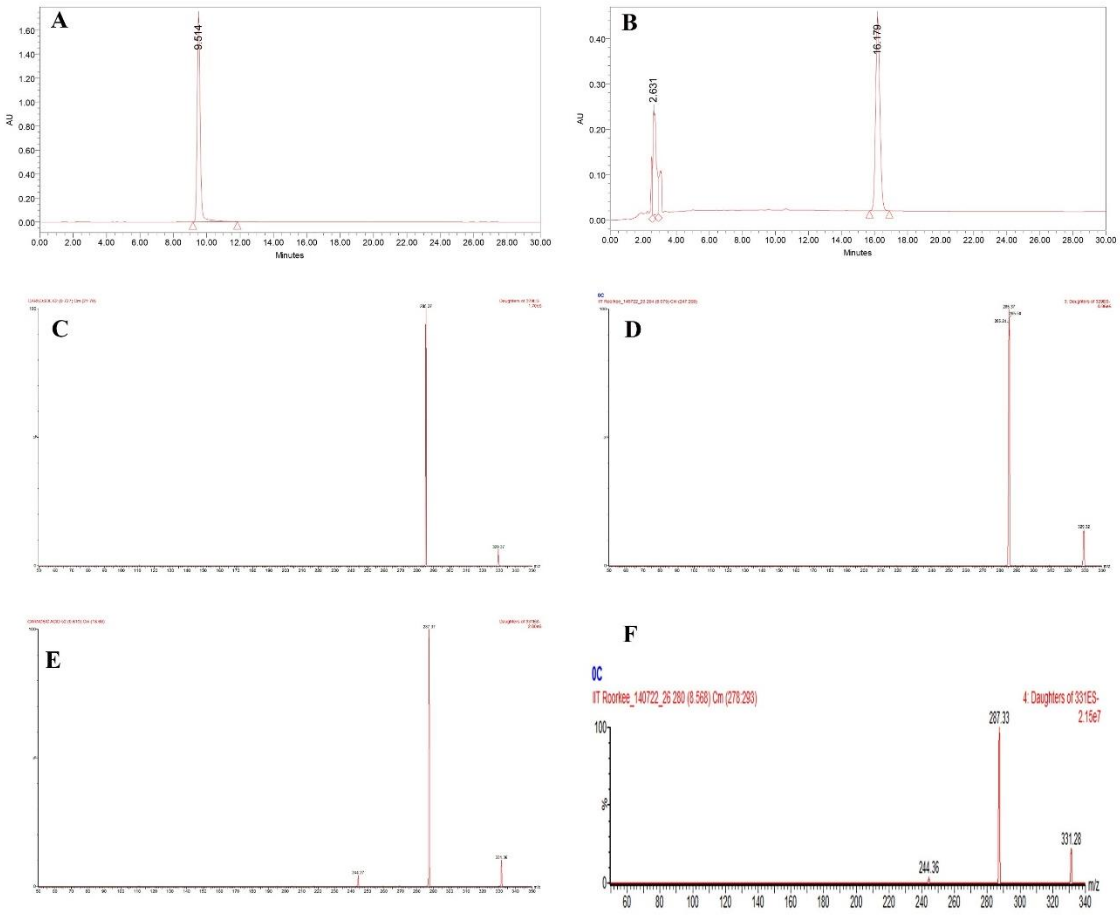


**Figure S2:** HPLC chromatograms and mass fragmentation patterns of CO and CA through LC-MS analysis. HPLC chromatograms of **A.** CO standard (9.5 min); **B**. CA standard (16.2 min); **C**. Mass fragmentation of CO standard (m/z: 329.37); **D**. Mass fragmentation of CO in leaf extracts of *S. officinalis* (m/z: 329.37). **E**. Mass fragmentation of CA standard (m/z: 331.28); **F**. Mass fragmentation of CA in leaf extracts of *S. officinalis* (m/z: 331.28).

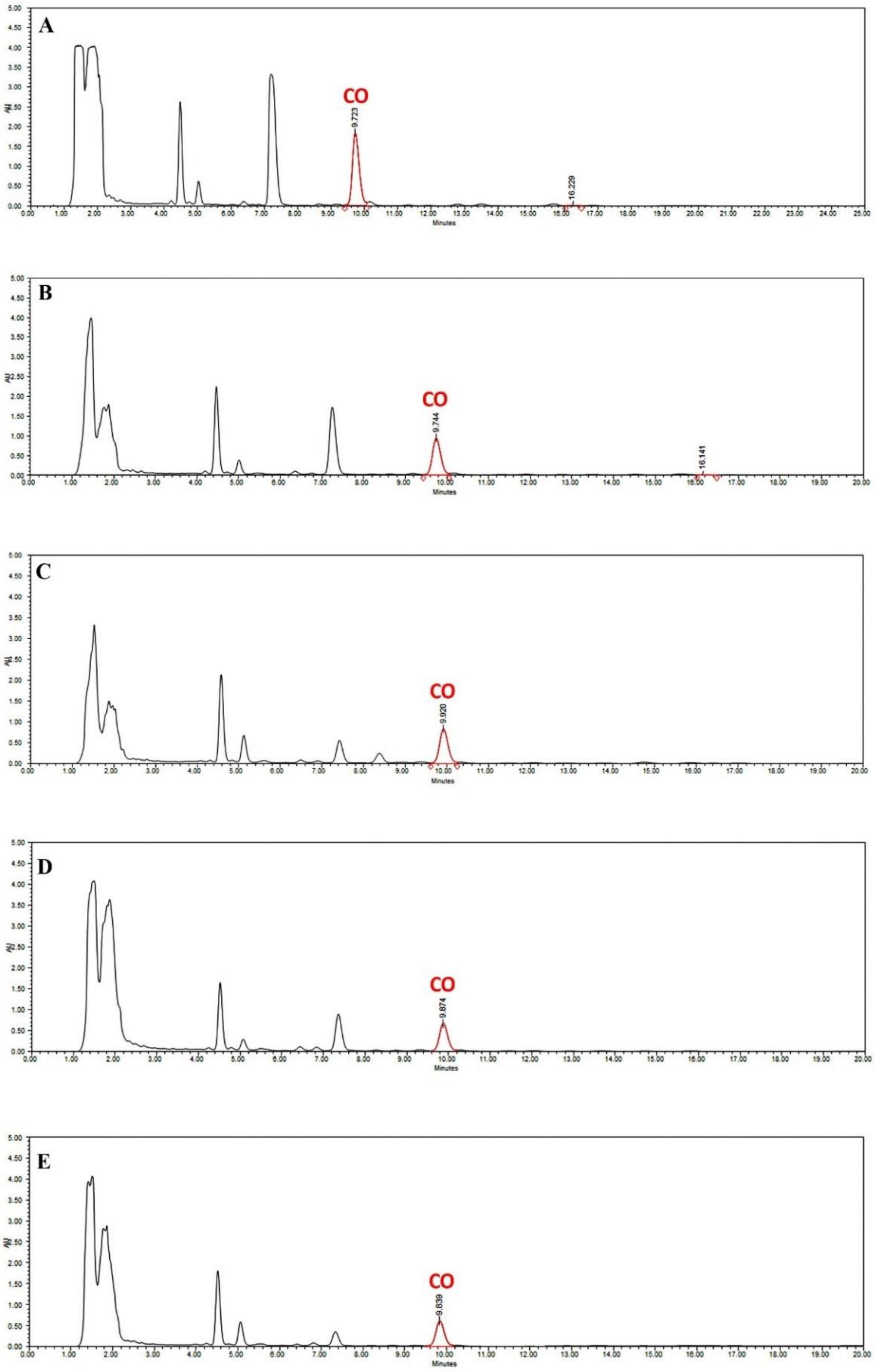


**Figure S3:** HPLC Chromatograms of different developmental stage leaf pairs extracts **A**. 1st pair; **B**. 2nd pair; **C**. 3rd pair, **D**. 4th pair; **E**. 5th pair.

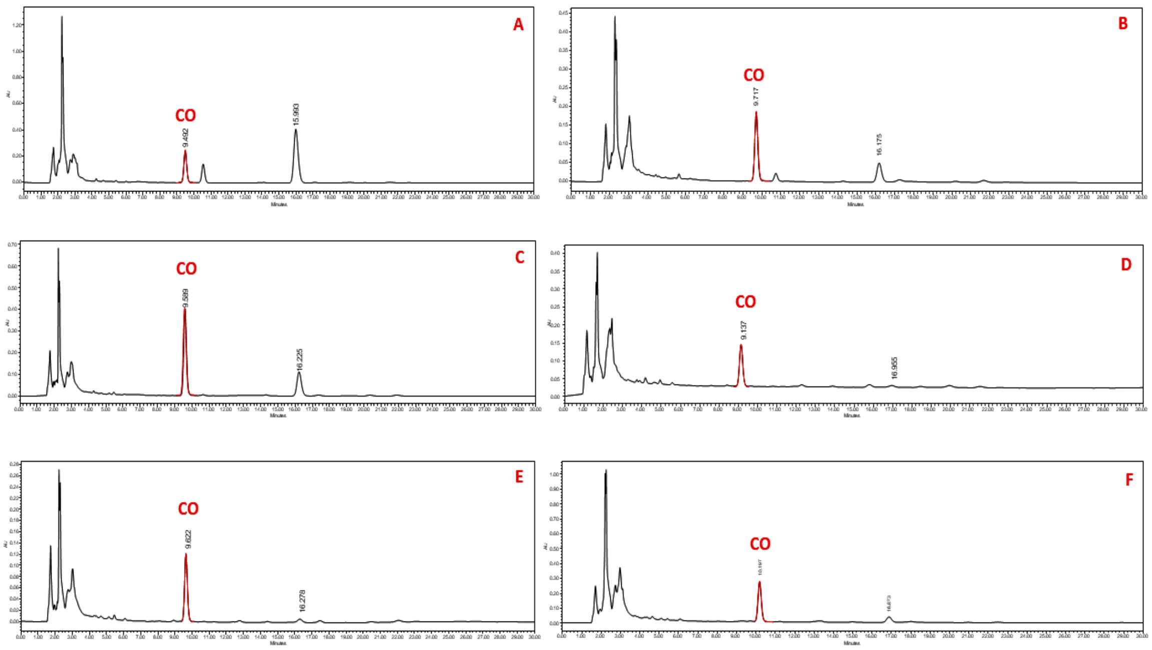


**Figure S4.** HPLC Chromatograms of sage leaf extracts post MeJA treatment. **A**. 0hrs, **B.** 2hrs, **C.** 6hrs, **D.** 12hrs, **E.** 24hrs, **F.** 48hrs.

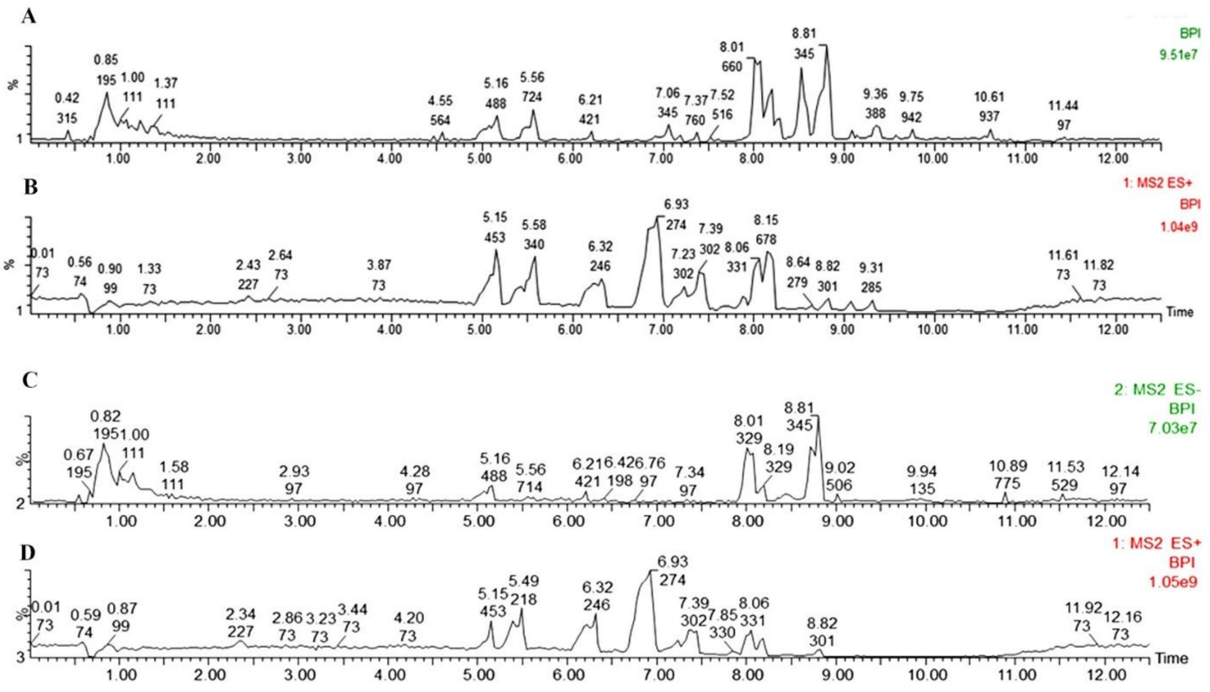


**Figure S5.** The total ion chromatograms (TIC) of *S. officinalis* leaf extracts infiltrated separately with *A. tumefaciens* harboring pBI121*:*EV and pBI121*::SmERF6* constructs. LC-MS was operated in both positive and negative ion modes to analyze the metabolite contents. **A**. Negative ion mode of pBI121*:*EV; **B**. Positive ion mode of pBI121*:*EV, **C.** Negative ion mode of pBI121*::SmERF6*, **D**. Positive ion mode of pBI121*::SmERF6*. The retention times of each detected metabolite and respective m/z values were provided in Table S3.

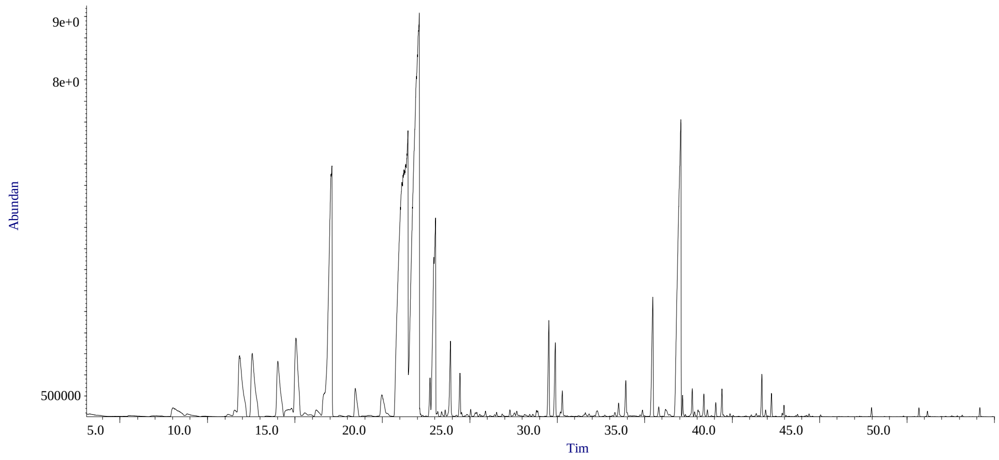


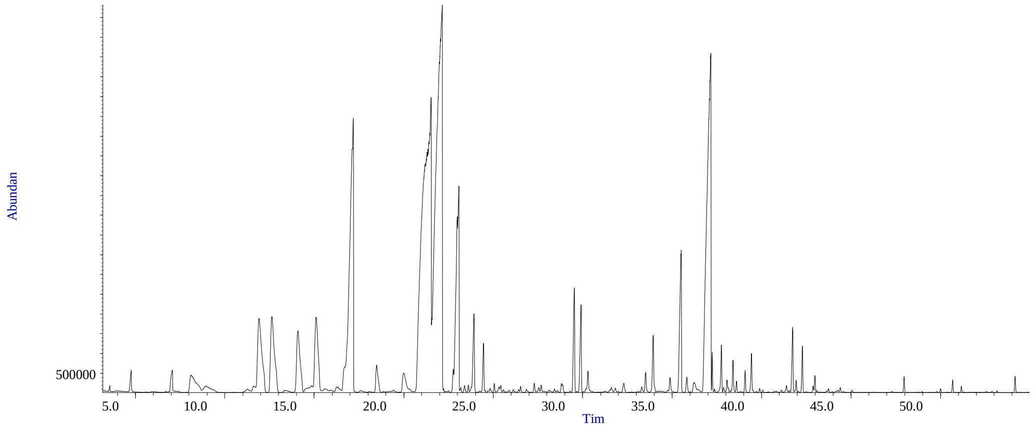


**Figure S6**. The GC-MS chromatograms of *S. officinalis* extracts infiltrated separately with *A. tumefaciens* harboring pBI121:EV and pBI121::*SmERF6* constructs. A. pBI121:EV, B. pBI121::*SmERF6*. The retention times of each detected metabolite and respective m/z values were provided in Table S4.

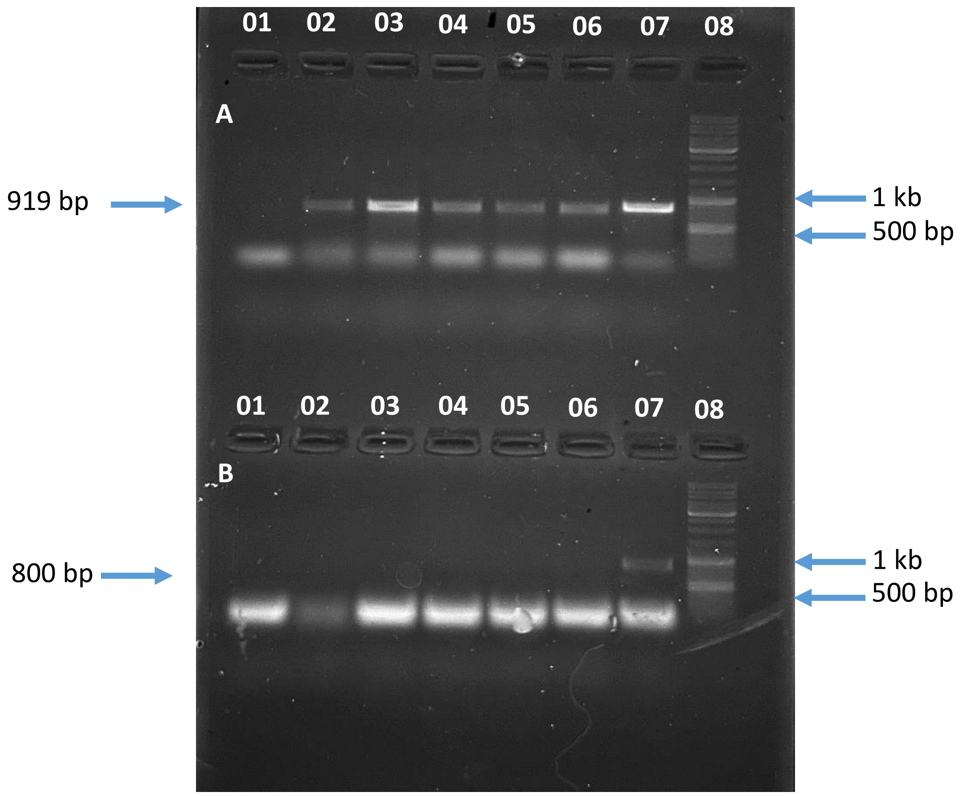


**Figure S7.** Screening of putative transgenic lines of S. *officinalis* transformed with *SmERF6*. A. lane shows the PCR amplification of integrated gene with *35S* forward and *SmERF6* reverse primers; 01: wild type, 02-06: putative transgenic lines, 07: positive control, 08: GeneRuler Ladder mix (Thermo scientific, USA). B. lane shows the PCR analysis to check the *Agrobacterium* contamination in putative transgenic lines by using *Chv* gene-specific primers; 01: wild type, 02-06: putative transgenic lines, 07: positive control, 08: GeneRuler Ladder mix (Thermo Scientific, USA).

| Genes | Primers |
|---|---|
| Real-time PCR primers | |
| *Elongation factor A (elfA)* | F: TTGTTGCCATTGACATCTTCACTT<br>R: CTCTCCCATGGCTGACATAACACT |
| *Geranylgeranyl pyrophosphate* (*GGPPS*) | F: GCTGCATTGTTAGAGGCCTC<br>R: CCCAACTCCTCCGAAGACTT |
| *Copalyl diphosphate synthase* (*CPS*) | F: AGATCACGGACTGGCTAGGA<br>R: TTTCAGCTGATGGCGAGTGT |
| *Kaurene Synthase Like* (*KSL*) | F: AGCACCGGGAAGAACTTGAG<br>R: GCAACGCGCAGTACATTTCT |
| *11-Hydroxy Ferruginol Synthase* (*HFS*) | F: CCCAACTTCGCCGACTACTT<br>R: CCGAAGTAAACATCCGCCCT |
| *xCYP76AK6/8* | F: TTGTGGCCAGCATTTTGAGC<br>R: ATACAATGGAGCATGCGGCT |
| *Sugiol Synthase* | F: CCCAACTTCGCCGACTACTT<br>R: CCCTCGATTAGAGCCAGCAG |
| *Salviol Synthase* | F: CAAGGATATGCGGCGAAACG<br>R: AGCCCATGCATTGACGATGA |
| Cloning Primers | |
| *SmERF6* | F: TCTAGA ATGATGGCAAATTCTGATGAGGGC<br>R: GAGCTC TCAAGAAGCCGGGTT |
| *SmERF6AS* | F: TCTAGA TCAAGAAGCCGGGTT<br>R: GAGCTC ATGATGGCAAATTCTG |
| *35s* | F: GCTCCTACAAATGCCATCA<br>R: GATAGTGGGATTGTGCGTCA |
| *Chv* | F: CGAAACGCTGTTCGGCCTGTGG<br>R: GTTCAGCAGGCCGGCATCCTGG |

**Table S1:** Primers used in the study

| Gene | Ipsort predicition | ChloroP | WOLFSPORT | Predotar |
|---|---|---|---|---|
| ***GGPPS*** | Chloroplast/mitochondrial | Chloroplast | Chloroplast | Elsewhere |
| ***CPS*** | Cytosolic | Elsewhere | Chloroplast | Plastidial |
| ***KSL*** | Chloroplast | Signal peptide | Chloroplast | Chloroplast |
| ***FS/HFS*** | Cytosolic | Chloroplast | Chloroplast | Endoplasmic reticulum |
| ***CYP76AK6*** | Chloroplast/mitochondrial | Elsewhere | Chloroplast | Endoplasmic reticulum |
| ***SmERF6*** | Nucleus | Nucleus | Nucleus | Nucleus |

**Table S2**: Sub-cellular localization of CO biosynthetic genes using various bioinformatics tools.

| S. No | Retention time | Compound | Mass (m/z) | Peak Abundance | |
|---|---|---|---|---|---|
| | | | | pBI121:*EV* | pBI121::*SmERF6* |
| 01 | 0.85 | Linalyl acetate | 196.2 | 1E08 ± 0.057 | 9E07 ± 0.20 |
| 02 | 1.79 | Humulene | 204.3 | 8E07 ± 0.057 | 7E07 ± 0.10 |
| 03 | 5.13 | Asiatic acid | 488.7 | 2E07 ± 0.057 | 3E07 ± 0.10 |
| 04 | 5.31 | Vanilic acid | 168.1 | 6E06 ± 0.057 | 1E07 ± 2.64 |
| 05 | 5.53 | Salvianolic acid | 494.4 | 1E06 ± 0.057 | 5E06 ± 0.55 |
| 06 | 5.95 | Linalool | 154.2 | 7E07 ± 0.057 | 2E08 ± 0.20 |
| 07 | 7.88 | Methyl rosmarinate | 374.3 | 2E07 ± 0.057 | 3E07 ± 0.23 |
| 08 | 7.92 | Rosmarinic acid | 360.3 | 1E06 ± 0.057 | 8E06 ± 0.57 |
| 09 | 7.95 | Sugiol | 300.4 | 6E05 ± 0.057 | 5E06 ± 0.20 |
| 10 | 8.13 | Ferruginol | 286.5 | 6E06 ± 0.057 | 1E07 ± 1.85 |
| 11 | 8.22 | Methoxy-carnosic acid | 346.5 | 9E05 ± 0.057 | 2E06 ± 1.0 |
| 12 | 8.22 | Rosmadial | 344.4 | 1E06 ± 0.057 | 5E06 ± 0.57 |
| 13 | 8.41 | Manool | 290.5 | 2E06 ± 0.057 | 2E06 ± 0.57 |
| 14 | 8.41 | Salviol | 302.5 | 7E06 ± 0.057 | 8E06 ± 0.68 |
| 15 | 8.52 | Viridiflorol | 222.37 | 3E06 ± 0.057 | 2E07 ± 0.57 |
| 16 | 8.61 | Caryophyllene | 204.35 | 1E07 ± 0.057 | 3E07 ± 0.15 |
| 17 | 8.78 | Epi rosmanol | 346.4 | 7E07 ± 0.057 | 9E07 ± 0.51 |
| 18 | 8.78 | Rosmanol | 346.4 | 3E06 ± 0.057 | 2E07 ± 0.57 |
| 19 | 7.82 | Ferrulic acid | 194.1 | 2E06 ± 0.057 | 1E07 ± 1.58 |
| 20 | 9.88 | Betulinic acid | 456.7 | 2E05 ± 0.057 | 3E06 ± 0.57 |

**Table S3**: List of compounds detected in leaf extracts of *S. officinalis* infiltrated with pBI121:EV and pBI121::*SmERF6* analyzed through LC-MS. The area of the compounds detected in respective samples were given in the table. The samples were run in triplicates and the data presented are average of three replicates.

| S. No | Retention time | Compound | Mass (m/z) | Peak abundance | |
|---|---|---|---|---|---|
| | | | | pBI121:EV | pBI121::*SmERF6* |
| 01 | 11.552 | 2-Thujene | 136.125 | 1E08 ± 0.03 | 1.9E08 ± 0.63 |
| 02 | 11.168 | (+)-3-Carene | 136.125 | 1.6E08 ± 0.03 | 1.4E08 ± 0.59 |
| 03 | 11.68 | α-Phellandrene | 136.125 | 2.3E08 ± 0.03 | 1.6E08 ± 0.58 |
| 04 | 11.841 | α-Pinene | 136.125 | 2.5E09 ± 0.03 | 3.1E09 ± 0.57 |
| 05 | 12.57 | Camphene | 136.125 | 2.4E09 ± 0.03 | 2.9E09 ± 0.68 |
| 06 | 14.027 | β-pinene | 136.125 | 1.9E09 ± 0.03 | 2.1E09 ± 0.58 |
| 07 | 15.061 | β-Myrcene | 136.125 | 2.6E09 ± 0.03 | 2.5E09 ± 0.87 |
| 08 | 16.207 | α-Terpinene | 136.230 | 3.8E08 ± 0.03 | 2.3E08 ± 0.62 |
| 09 | 16.691 | p-Cymene | 134.11 | 4.7E08 ± 0.03 | 4.7E08 ± 0.72 |
| 10 | 17.119 | Eucalyptol | 154.136 | 8.3E09 ± 0.03 | 9.8E09 ± 0.59 |
| 11 | 18.454 | γ-Terpinene | 136.125 | 8E08 ± 0.03 | 6.3E08 ± 0.85 |
| 12 | 19.972 | Terpinolene | 136.230 | 9.3E08 ± 0.03 | 6.7E08 ± 0.80 |
| 13 | 21.112 | Thujone | 152.12 | 4.4E10 ± 0.03 | 5.2E10 ± 0.66 |
| 14 | 22.725 | 3-Thujanol | 154.136 | 4.8E08 ± 0.03 | 2.6E08 ± 0.49 |
| 15 | 22.942 | (+)-2-Bornanone | 152.12 | 2.5E09 ± 0.03 | 2.9E09 ± 0.44 |
| 16 | 23.387 | Sabinone | 150.104 | 1.7E08 ± 0.03 | 9.9E07 ± 0.93 |
| 17 | 23.893 | endo-Borneol | 154.136 | 1.1E09 ± 0.03 | 1.1E09 ± 0.72 |
| 18 | 24.432 | Terpinen-4-ol | 154.136 | 5.1E08 ± 0.03 | 5.5E08 ± 0.80 |
| 19 | 25.044 | α-Terpineol | 154.136 | 1.5E08 ± 0.03 | 9.4E07 ± 1.14 |
| 20 | 28.12 | Carvenone | 152.12 | 7.9E07 ± 0.03 | 5.6E07 ± 0.77 |
| 21 | 29.516 | Bornyl acetate | 196.146 | 1.1E09 ± 0.03 | 1.1E09 ± 0.54 |
| 22 | 29.649 | Thymol | 150.104 | 3.8E08 ± 0.03 | 3.2E08 ± 0.44 |
| 23 | 29.888 | Sabinol isovalerate | 236.178 | 9.3E08 ± 0.03 | 1.1E09 ± 0.6 |
| 24 | 32.285 | α-Cubebene | 204.188 | 2.3E08 ± 0.03 | 1.7E08 ± 0.54 |
| 25 | 32.719 | Eugenol | 164.084 | 1.2E08 ± 0.03 | 1.2E07 ± 0.18 |
| 26 | 33.297 | Ylangene | 204.188 | 1.3E08 ± 0.03 | 8.3E07 ± 0.72 |
| 27 | 33.509 | Copaene | 204.188 | 2.7E08 ± 0.03 | 4.8E08 ± 0.81 |
| 28 | 33.92 | (-)-β-Bourbonene | 204.188 | 5.6E08 ± 0.03 | 7.7E08 ± 0.77 |
| 29 | 34.871 | Caryophyllene | 204.188 | 1.8E09 ± 0.03 | 2.6E09 ± 0.35 |
| 30 | 36.189 | Aromandendrene | 204.188 | 3.1E08 ± 0.03 | 2.9E08 ± 0.84 |
| 31 | 36.44 | γ-Muurolene | 204.188 | 1.2E08 ± 0.03 | 5.7E08 ± 1.18 |
| 32 | 37.168 | Alloaromadendrene | 204.188 | 2.2E08 ± 0.03 | 2.9E08 ± 0.89 |
| 33 | 38.586 | α -Muurolene | 204.188 | 3.3E08 ± 0.03 | 3.1E08 ± 1.04 |
| 34 | 39.699 | Cubenene | 204.188 | 7.5E07 ± 0.03 | 1.8E07 ± 0.39 |
| 35 | 40.06 | α -Calacorene | 200.157 | 8.4E07 ± 0.03 | 3.3E07 ± 0.60 |
| 36 | 41.384 | Caryophyllene oxide | 220.183 | 1.1E08 ± 0.03 | 9.7E07 ± 1.04 |
| 37 | 41.701 | Viridiflorol | 222.198 | 4.5E08 ± 0.03 | 7.3E08 ± 0.41 |
| 38 | 41.918 | Humulene epoxide I | 220.183 | 1.3E08 ± 0.03 | 1.2E08 ± 0.80 |

**Table S4**: List of VOCs detected in leaf extracts of *S. officinalis* infiltrated with pBI121:EV and pBI121::*SmERF6* analyzed through GC-MS. The samples were run in triplicates and the data presented are average of three replicates.

| | |
|---|---|
| Total Number of treated plants | 347 |
| Total number of plants survived after treatment | 115 |
| Total number of plants screened for transgene integration | 115 |
| Total number of confirmed lines | 05 |
| Total number of plants survived | 03 |
| Transformation efficiency | 4.3% |

**Table S5**: Efficiency *of in planta Agrobacterium* mediated genetic transformation in *S. officinalis*
